# Gut-derived bacterial flagellin induces beta-cell inflammation and dysfunction

**DOI:** 10.1101/2021.10.07.463317

**Authors:** Torsten P.M. Scheithauer, Hilde Herrema, Hongbing Yu, Guido J. Bakker, Maaike Winkelmeijer, Galina Soukhatcheva, Derek Dai, Caixia Ma, Stefan R. Havik, Manon Balvers, Mark Davids, Abraham S. Meijnikman, Ömrüm Aydin, Bert-Jan H. van den Born, Marc G. Besselink, Olivier R. Busch, Maurits de Brauw, Arnold van de Laar, Clara Belzer, Martin Stahl, Willem M. de Vos, Bruce A. Vallance, Max Nieuwdorp, C. Bruce Verchere, Daniël H. van Raalte

## Abstract

**Objective:** Hyperglycemia and type 2 diabetes (T2D) are caused by failure of pancreatic beta cells. The role of the gut microbiota in T2D has been studied but causal links remain enigmatic.

**Design:** Obese individuals with or without T2D were included from two independent Dutch cohorts. Human data was translated *in vitro* and *in vivo* by using pancreatic islets from C57BL6/J mice and by injecting flagellin into obese mice.

**Results:** Flagellin is part of the bacterial locomotor appendage flagellum, present on gut bacteria including Enterobacteriaceae, which we show to be more abundant in the gut of individuals with T2D. Subsequently, flagellin induces a pro-inflammatory response in pancreatic islets mediated by the Toll-like receptor (TLR)-5 expressed on resident islet macrophages. This inflammatory response associated with beta-cell dysfunction, characterized by reduced insulin gene expression, impaired proinsulin processing and stress-induced insulin hypersecretion *in vitro* and *in vivo* in mice.

**Conclusion:** We postulate that increased systemically disseminated flagellin in T2D is a contributing factor to beta cell failure in time and represents a novel therapeutic target.

**Graphical abstract:** 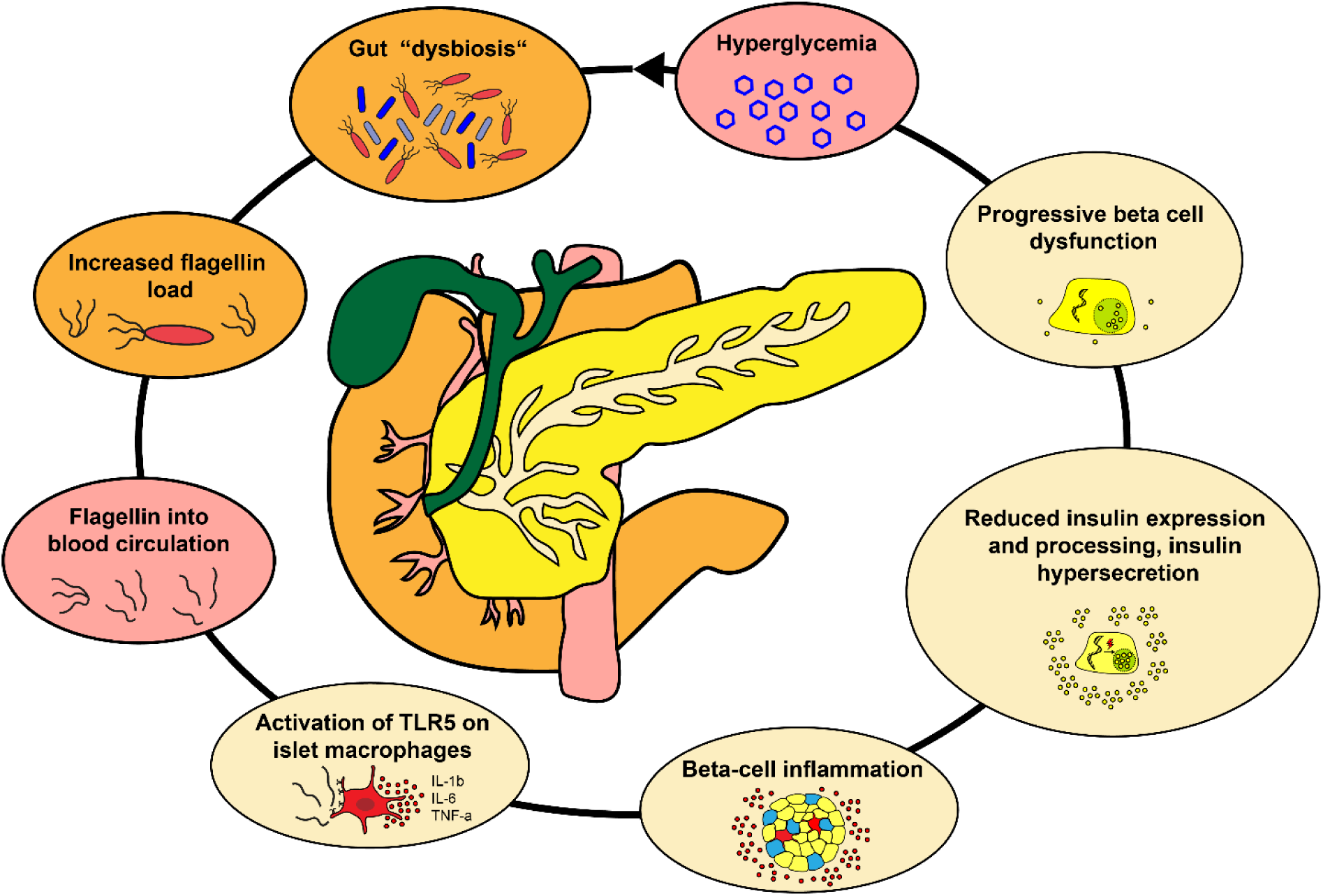

## Introduction

While obesity is linked to insulin resistance, it is failure of pancreatic beta-cells that drives hyperglycemia and subsequent type 2 diabetes (T2D) (Weyer et al., 1999). Although in later stages of T2D insulin secretory rates are lowered, prior to the diagnosis and in earlier phases of the disease, insulin secretion is actually increased (Defronzo, 2009). Insulin hypersecretion, particularly in the fasted state, is considered harmful as it associates with impaired proinsulin processing, insulin secretory stress and depletion of intracellular insulin stores (Pories and Dohm, 2012), further promoting obesity and T2D development (Mehran et al., 2012; Tricò et al., 2018; Weyer et al., 2000). Drivers of hyperinsulinemia are still elusive, but could relate to islet-exposure to excessive nutrients such as carbohydrates and lipids (Erion and Corkey, 2018), as well as a chronic low-grade inflammatory response known to be present in beta cells of people with T2D. In this regard, an influx of pro-inflammatory macrophages in islets of people with T2D has been noted (Donath and Shoelson, 2011; Marchetti, 2016). These macrophages produce pro-inflammatory cytokines such as interleukin (IL)-1β and IL-6, which have been associated with insulin hypersecretion (Ellingsgaard et al., 2011; Hajmrle et al., 2016) and beta-cell failure (Donath et al., 2009; Donath and Shoelson, 2011). The triggers that ignite beta-cell inflammation in T2D remain presently unknown.

A recent player in the field of glucose metabolism is the intestinal microbiota. Several cohort (Le Chatelier et al., 2013) and intervention (Kootte et al., 2017) studies have shown an association between gut microbiota composition and T2D incidence (Gurung et al., 2020). People with obesity and T2D often have lower microbial diversity, while showing increased abundance of potentially pathogenic gram-negative bacteria, including Proteobacteria (Ouchi et al., 2011). Mechanistic studies have linked metabolites produced by the gut microbiota to impaired glucose metabolism and a pro-inflammatory state (Herrema and Niess, 2020). In addition to microbial metabolites, structural components of gram-negative bacteria, such as lipopolysaccharide (LPS), a cell-wall component, and flagellin, part of the bacterial locomotor appendage flagellum, may systemically disseminate in people with T2D (Gomes et al., 2017). These bacterial components activate pro-inflammatory pathways by binding to pattern-recognition receptors (PRRs), including Toll-like receptors (TLRs), expressed on epithelial cells and cells of the innate immune system (Scheithauer et al., 2020).

Here, we provide evidence for a novel pathway in which exaggerated systemic dissemination of gut-derived flagellin in T2D induces a pro-inflammatory state in beta-cells. This inflammatory response is mediated by flagellin-mediated activation of TLR5 expressed on resident islet macrophages. Functionally, the inflammatory response associates with impaired insulin gene expression and proinsulin processing, while inducing hyperinsulinemia. Collectively, these processes markedly reduce insulin stores, which potentially contribute to beta-cell failure over time.

## Results

### Fecal Enterobacter cloacae abundance is associated with hyperglycemia in humans

To investigate the link between beta-cell dysfunction and altered gut microbiota, we analyzed fecal samples for microbiota composition using 16S rRNA sequencing in participants enrolled in the Healthy Life in an Urban Setting (HELIUS) study, a prospective cohort study of the six largest ethnic groups living in Amsterdam, The Netherlands (Deschasaux et al., 2018; Snijder et al., 2017). To prevent confounding effects of ethnic differences on gut microbiota composition (Deschasaux et al., 2018), we analyzed the samples of the 803 Dutch origin participants (**Table S1**). We observed increased abundance of Gram-negative Enterobacteriaceae in people with T2D as compared to normoglycemic controls (**Figure 1A**, **Table S2**), confirming a previous report where Enterobacteriaceae were increased in people with T2D (Qin et al., 2012). We randomly selected 100 people with T2D and compared them to 50 age-, sex- and BMI-matched normoglycemic controls also recruited within the HELIUS cohort (**Table S3**). We confirmed an enrichment of Enterobacteriaceae in people with T2D using quantitative polymerase chain reaction (qPCR) (**Figure 1B**). Furthermore, we observed a positive relation with the long-term glucose marker hemoglobin A1c (HbA1c) and Enterobacteriaceae abundance (**Figure 1C**).

**Figure 1.**
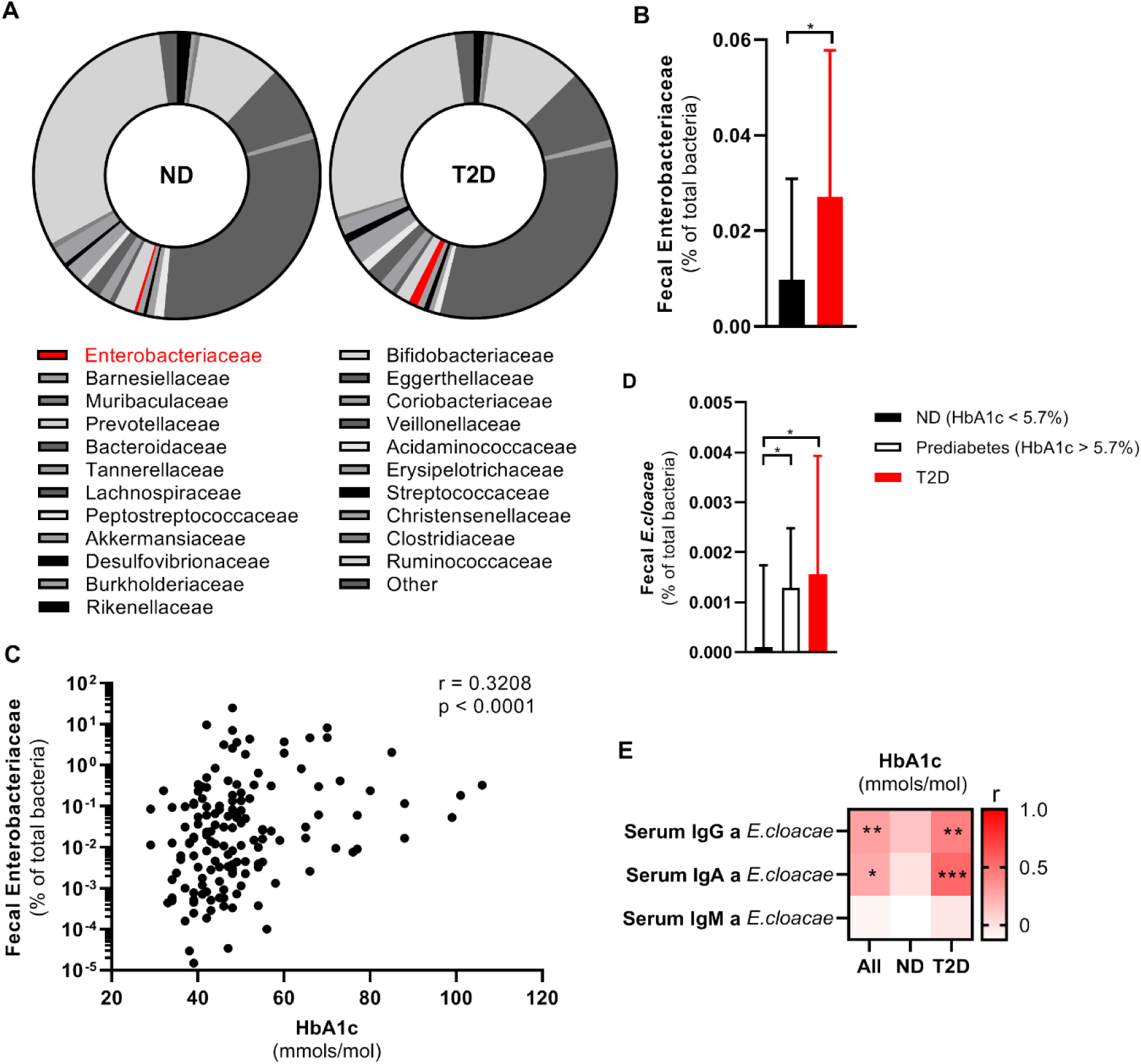
Fecal Enterobacteriaceae is associated with a disturbed glucose tolerance in humans from the HELIUS cohort. (A) Fecal microbiota composition of people with or without T2D measured via 16S rRNA sequencing (Dutch origin participants, N = 803, % abundance, median is shown). (B) Fecal Enterobacteriaceae (qPCR, normalized to total fecal bacterial DNA) is increased in individuals with T2D compared to age-BMI-sex matched healthy controls (N = 150, median with 95% CI). (C) Fecal Enterobacteriacaee (qPCR, normalized to total fecal bacteria) positively correlates with the long-term glucose marker HbA1c (N = 150). (D) Fecal *Enterobacter cloacae* (qPCR, normalized to total fecal bacteria) is increased in prediabetes and T2D (N = 150, median with 95% CI). (E) Correlation analysis of serum antibodies against *E. cloacae* and HbA1c (N = 80). Mann Whitney test (B, D) and Spearman correlation (C, E); *p<0.05, **p<0.01, ***p<0.001. Abbreviations: ND, no diabetes; T2D, type 2 diabetes; HbA1c, Glycated hemoglobin; Ig, immunoglobulin; CI, confidence interval.

*Enterobacter cloacae (E. cloacae)*, a prominent member of the family of Enterobacteriaceae, was previously shown to be associated with impaired glucose tolerance in humans and mice (Fei and Zhao, 2013; Keskitalo et al., 2018). In line with these studies, in our cohort, levels of *E. cloacae* directly increased with deterioration of glucose tolerance (**Figure 1D**). Further, fecal abundance of *E. cloacae* also positively correlated with HbA1c (**Figure S1A**). Thus, as a proof-of-concept, we selected *E. cloacae* for subsequent experiments although we acknowledge that other bacteria of the family Enterobacteriaceae may also associate with glucose (dys)metabolism.

### An immune response against Enterobacter cloacae is associated with hyperglycemia in people with type 2 diabetes

An appropriate immune response to opportunistic bacteria is necessary to prevent inflammation (Cullender et al., 2013b). To assess whether there was a systemic immune response to *E. cloacae*, we assessed plasma antibody levels. We observed a numerical increase in IgG titers against *E. cloacae* in T2D, but otherwise no significant difference between the matched groups with respect to antibodies was noted (**Figure S1B**). However, there was a positive correlation between HbA1c levels and systemic IgG and IgA against *E. cloacae* (**Figure 1E**), particularly in people with T2D. Further, there was a significant positive correlation between fecal abundance of Enterobacteriaceae and plasma IgG against *E. cloacae* (**Figure S1C**). This is suggestive of an immune response against systemically disseminated bacterial components of *E. cloacae*.

### Enterobacter cloacae induces beta-cell inflammation and dysfunction in vitro

Given the link between beta-cell driven hyperglycemia and fecal presence of *E. cloacae* as well as systemic antibodies against *E. cloacae*, we questioned if *E. cloacae* would be able to alter pancreatic beta-cell function. We isolated pancreatic islets from C57BL6/J mice fed a conventional chow diet. Islets were co-incubated with 10^6^ colony forming units (CFUs) per mL of heat-inactivated *E. cloacae* or vehicle for 72 hours (**Figure 2A-G**). We found that beta-cells exposed to heat-inactivated *E. cloacae* had lower expression of genes involved in insulin production, including the key transcription factors pancreatic duodenal homeobox 1 (*PDX1*) and *MafA* (**Figure 2A**). This lowered expression coincided with a higher inflammatory tone (**Figure 2B** and **2C**), including upregulation of pro-inflammatory cytokines (*IL-1β*, *IL-6* and tumor necrosis factor (*TNF-α*), the *NLRP3* inflammasome, the macrophage marker *F4/80*, and *TLR2*. Interestingly, increased TLR2 expression was previously reported in pancreatic islets of people with diabetes (Ji et al., 2019). Heat-inactivated *E. cloacae* did not affect cell viability since ATP content was not altered (**Figure 2D**) (Iyer et al., 2009). Incubation with heat-inactivated *E. cloacae* also had functional consequences for beta cells. As such, insulin content was markedly reduced after 72 hours of incubation with *E. cloacae* (**Figure 2E**). In addition, both during low- and high ambient glucose concentrations, beta cells treated with heat-inactivated *E. cloacae* hypersecreted insulin (**Figure 2F**). Lastly, *E. cloacae* treatment increased proinsulin secretion and content as well as proinsulin/insulin ratios, indicating disturbed proinsulin processing (**Figure 2G**). We observed similar data in human islets, where heat-inactivated *E. cloacae* lowered *MafA* expression, increased secreted IL-6, and tended to reduce insulin content (**Figure S2A-E**). Thus, the profile of increased inflammation, reduced insulin gene expression, impaired proinsulin processing and insulin hypersecretion likely contributes to the detrimental reduction in beta-cell insulin content.

**Figure 2.**
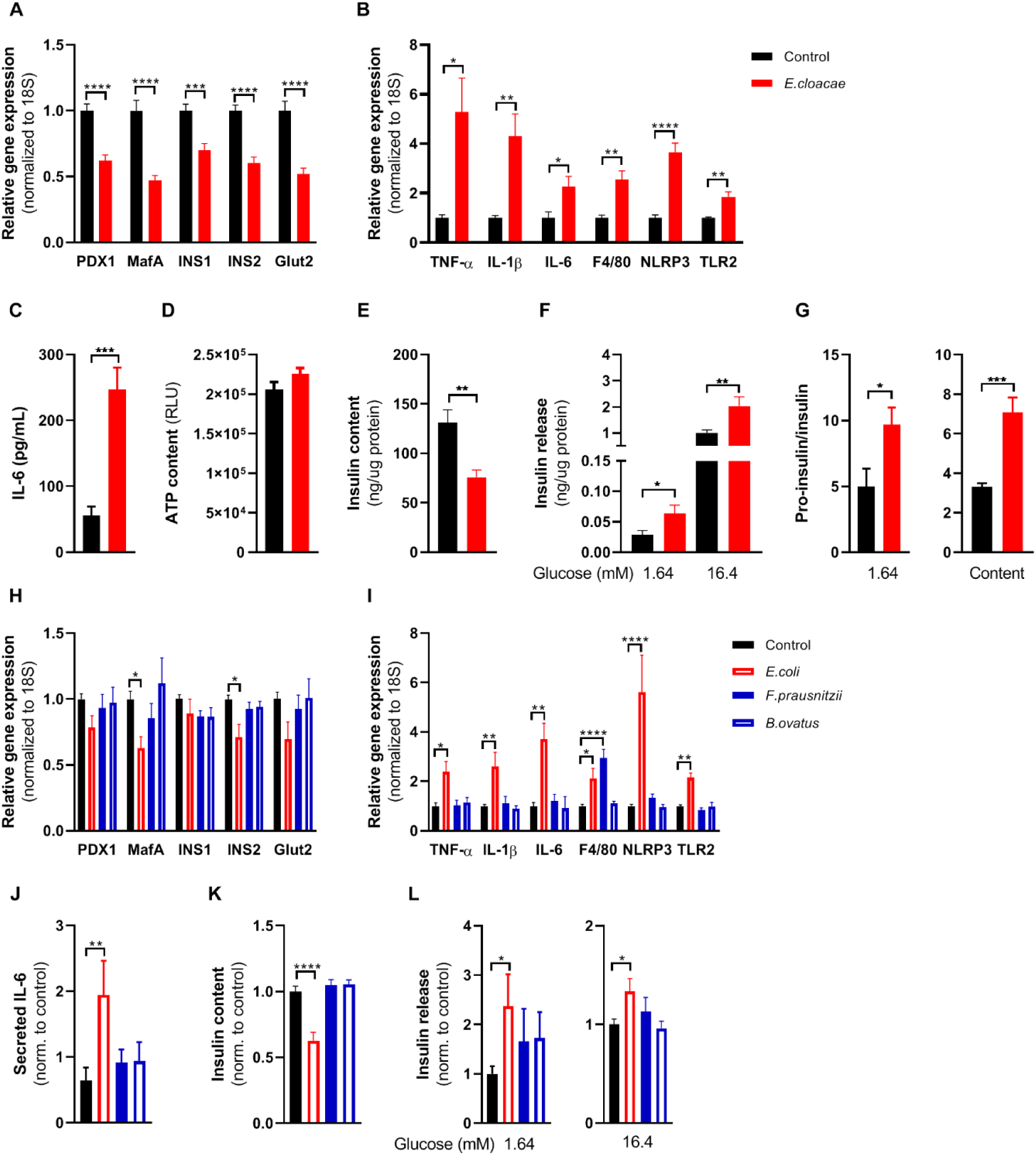
Opportunistic pathogens, but not beneficial bacteria induce beta cell inflammation and dysfunction. Freshly isolated pancreatic islets from healthy C57BL6J mice were treated with heat-inactivated bacteria for 72h (1E6 colony forming units/mL). (A) *Enterobacter cloacae* (E. cloacae) reduces expression of beta-cell genes. (B) *E. cloacae* increases expression of inflammatory genes in islets. (C) *E. cloacae* increases IL-6 secretion by islets. (D) *E. cloacae* does not reduce ATP content in islets. (E) *E. cloacae* reduces insulin content in islets. (F) *E. cloacae* increases insulin secretion from islets during low- and high glucose conditions, denoting insulin hypersecretion. (G) *E. cloacae* increases the ratio between secreted pro-insulin and insulin, as well as the islet content of pro-insulin relative to insulin, indicating impaired insulin processing. (H) *Escherichia coli*, but not *Faecalibacterium prausnitzii and Bacteroides ovatus*, reduces expression of beta-cell genes (I) *E. coli*, but not *F. prausnitzii and B. ovatus* increases expression of inflammatory genes in islets. (J) *E. coli*, but not *F. prausnitzii and B. ovatus*, induces the release of IL-6 from islets into the media. (K) *E. coli*, but not *F. prausnitzii and B. ovatus* reduces insulin content in islets. (L) *E. coli*, but not *F. prausnitzii and B. ovatus* induces insulin hypersecretion versus controls at both low- and high-glucose conditions. Data shown are mean ± SEM. Unpaired t-test (A, B, C, D, G, H, I, J) and Mann Whitney test (E, F, K, L) was used. Significance level: *p<0.05, **p<0.01, ***p<0.001, ****p<0.0001 (mean ± SEM, 3 representative experiments per panel). Gene expression was normalized using *18s* as a housekeeping gene. Panels J-L were normalized to the control samples since the experiments were performed independently (each bacterium on different days). Abbreviations: PDX1, pancreatic and duodenal homeobox 1; INS1 and INS2, insulin 1 and 2; NLRP3, NACHT, LRR and PYD domains-containing protein 3; TNF-α, tumor necrosis factor-alpha; IL-1β, Interleukin 1 beta; IL-6, Interleukin 6; TLR2, Toll-like receptor 2; RLU, Relative light unit.

### Opportunistic pathogens, but not beneficial bacteria, induce beta-cell inflammation and dysfunction

Next, to address whether the *E. cloacae*-mediated effects were specific to this bacterial species, we repeated the experiments with *Escherichia coli (E. coli)*, another Gram-negative bacterium from the group Enterobacteriaceae (Amar et al., 2011a). *E. coli* was also increased in T2D participants of the HELIUS study (**Figure S1D**). In line with our *E. cloacae* findings*, E. coli* reduced expression of genes regulating beta-cell maturation and function (**Figure 2H**) and induced an inflammatory response with increased expression of *IL-1β*, *IL-6*, *TNF-α*, the *NLRP3* inflammasome, *F4/80* and *TLR2* (**Figure 2I**), and IL-6 protein secretion (**Figure 2J**). *E. coli* also reduced cellular insulin content (**Figure 2K**) and increased insulin secretion (**Figure 2L**).

In order to rule out an effect of bacterial co-incubation *per se*, we investigated the effects of two bacteria that have been identified as beneficial for the host. These included the Gram-positive *Faecalibacterium prausnitzii* (Qin et al., 2012) and Gram-negative *Bacteroides ovatus* (Zhang et al., 2014), the abundance of which was decreased in people with T2D (**Figure S1E** and **S1F**). In contrast to *E. coli* and *E. cloacae*, *F. prausnitzii* and *B. ovatus* did not affect islet inflammation, insulin content or insulin secretion (**Figure 2H-L**). These data indicate that only a subset of bacteria induce an inflammatory response and beta-cell dysfunction. Based on previous mouse data linking *E. cloacae* to impaired glucose tolerance (Fei and Zhao, 2013), we decided to further scrutinize the effect of this bacterium on beta-cell function as proof-of-concept.

### Toll-like receptor-2 and Toll-like receptor-4 deletion do not protect against Enterobacter cloacae-induced beta-cell inflammation and dysfunction

TLR2 and TLR4 are involved in beta-cell replication (Ji et al., 2019) and have been proposed as two key PRR’s that mediate the inflammatory response induced by endogenous and exogenous molecules, the latter including bacterial components such as LPS (Takeuchi et al., 1999). In addition, TLR2 and TLR4 are expressed by pancreatic islet cells (Akira and Takeda, 2004; Giarratana et al., 2004; Wen et al., 2004). Therefore, we isolated islets from TLR2 and TLR4 knock out mice and incubated them with *E. cloacae* (**Figure S3**). Despite the absence of TLR2*, E. cloacae* reduced insulin gene expression including *MafA* (**Figure S3A**), increased expression of pro-inflammatory cytokines (**Figure S3B**), increased secreted IL-6 (**Figure S3C**), and reduced insulin content (**Figure S3D**).

Similarly, TLR4-deficient islet cells were not protected from the effects of *E. cloacae*, as the expression of beta-cell genes including *INS2* was still reduced (**Figure S3F**). Regarding inflammation, while the expression of *IL-1β, NLRP3* inflammasome and *F4/80* was similarly increased by *E. cloacae* in both TLR4 knock out and WT islets, *E. cloacae* incubation did not increase *IL-6* expression and secretion (**Figure S3G-H**). Insulin content was also reduced by *E. cloacae* incubation (**Figure S3I and S3J**). Therefore, we concluded that PRRs other than TLR2 and TLR4 likely play roles in *E. cloacae*-induced beta-cell inflammation and dysfunction.

### Toll-like receptor-5 deletion protects against Enterobacter cloacae-induced beta-cell inflammation and dysfunction

Several members of Enterobacteriaceae, including *E. cloacae*, express flagellins as both virulence and motility factors (De Maayer and Cowan, 2016). Bacterial flagellin is mainly recognized by TLR5 (Yoon et al., 2012), which is expressed by various cell types including epithelial cells and monocytes. We measured the effects of *E. cloacae* in islets from TLR5 knock out mice. TLR5 deletion partially protected islets from beta-cell dysfunction with preserved expression of insulin genes (**Figure 3A**). TLR5 deficiency did not reduce the effects of *E. cloacae* on expression of pro-inflammatory cytokines (**Figure 3B**), although it did reduce secretion of IL-6 as compared to WT islets (**Figure 3C**). In addition, in TLR5 knock out islets, *E. cloacae* did not reduce insulin content (**Figure 3D**), while similar insulin secretion rates were observed versus WT islets (**Figure 3E**).

**Figure 3.**
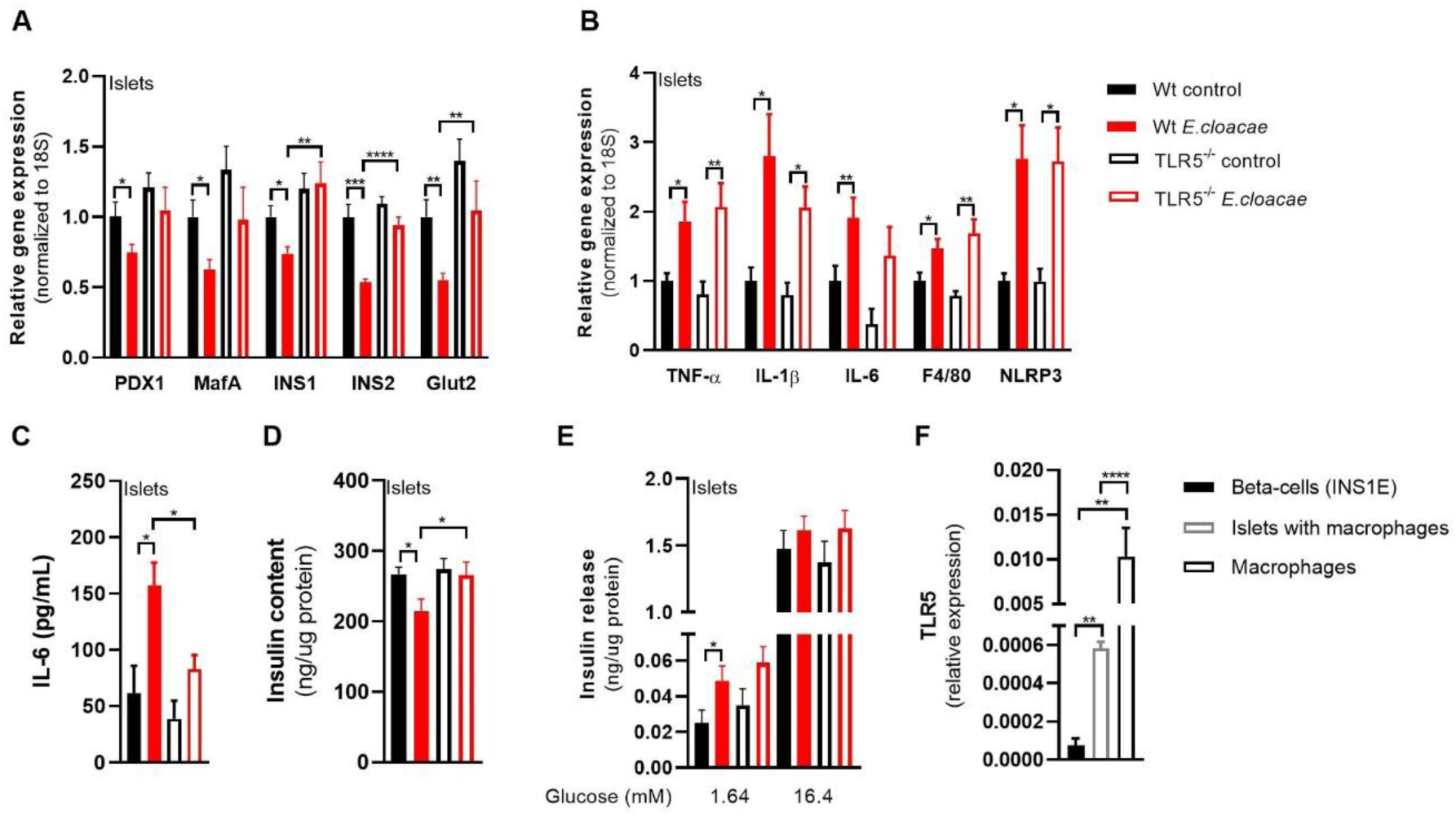
TLR5 is mediating beta cell dysfunction in pancreatic islets. Freshly isolated pancreatic islets from C57BL6J TLR5^-/-^ mice (10 weeks old) were treated with heat-inactivated *Enterobacter cloacae* (1E6 CFUs/mL) for 72h. (A) *E. cloacae* reduces beta-cell gene expression. TLR5 knock out protects from *E. cloacae*-induced insulin and Glut2 expression loss. (B) *E. cloacae* increases beta-cell inflammation. TLR5 knock out does not prevent the effect of *E. cloacae* on pancreatic islet inflammation. (C) *E. cloacae* increases beta-cell IL-6 secretion. TLR5 knock out partially protects from *E. cloacae* induced IL-6 secretion. (D) *E. cloacae* reduces beta-cell insulin content. TLR5 knock out protects from *E. cloacae*-induced insulin content loss. (E) *E. cloacae* induces insulin hypersecretion at low glucose condition in wild-type islets, which is not different in TLR5 knock out islets. (F) Clonal beta cells express very low levels of TLR5 compared to pancreatic islets and macrophages. Data shown are mean ± SEM. Unpaired t-test was used for statistical analysis (A-C, F) or Mann-Whitney test (D, E). Significance level: *p<0.05, **p<0.01, ***p<0.001, ****p<0.0001 (mean ± SEM, 3 representative experiments per panel). Abbreviations: PDX1, pancreatic and duodenal homeobox 1; INS1 and INS2, insulin 1 and 2; NLRP3, NACHT, LRR and PYD domains-containing protein 3; TNF-α, tumor necrosis factor-alpha; IL-1β, Interleukin 1 beta; IL-6, Interleukin 6; TLR2, Toll-like receptor 2; TLR5, Toll-like receptor 5.

To further assess the role of TLR5 in mediating the effects of *E. cloacae*, we co-incubated WT primary mouse islets with *E. cloacae* with or without the TLR5 inhibitor TH1020. As TH1020 proved toxic to cells after prolonged incubation, we studied the islets after 6 hours of treatment. In line with TLR5 knock out islets, TH1020 reduced the effects of *E. cloacae* on beta-cell gene expression (partial preservation of *PDX1* and *MafA* expression) and partially offset the *E. cloacae*-induced expression of *IL-1β, IL-6* and *NLRP3* (**Figure S4A-B**). Due to the toxic effects of TH1020, particularly on GLUT2 expression, we did not perform glucose stimulated insulin secretion (GSIS).

### Macrophages mediate Enterobacter cloacae-induced beta cell inflammation and dysfunction via TLR5 activation

As TLR5 is barely expressed in beta cells (**Figure 3F**), we concluded that other islet associated cells in the pancreas, particularly resident islet macrophages, could mediate the observed effects of TLR5 activation. Indeed, macrophages as well as pancreatic islets containing macrophages had higher TLR5 expression than pure beta cells (**Figure 3F**). Macrophages are important for pancreatic islet physiology (Nackiewicz et al., 2020), but can induce beta-cell dysfunction when activated towards a pro-inflammatory phenotype (Ying et al., 2019). We thus depleted macrophages from murine pancreatic islets using clodronate-liposomes (Nackiewicz et al., 2014), followed by *E. cloacae* incubation (**Figure 4A-E**). Reduction of islet macrophages resulted in the maintenance of beta-cell gene transcription following *E. cloacae* treatment (**Figure 4A**). Further, expression of pro-inflammatory cytokines including *IL-1β* and *NLRP3* was reduced (**Figure 4B**). Particularly, IL-6 secretion was almost abolished (**Figure 4C**). Insulin content (**Figure 4D**) and insulin secretion (**Figure 4E**) were not affected by *E. cloacae* in islets lacking macrophages. In line with a role for islet resident macrophages, we found that pure beta cells (INS1E clonal cell line) did not show inflammation or beta-cell dysfunction following *E. cloacae* treatment (**Figure S5A-D**). Further, human islet organoids that consist of pure human endocrine cells, did not show signs of beta-cell dysfunction upon *E. cloacae* treatment (**Figure S5E-F**). Further evidence for the mediating role of macrophages, *E. coli* and *E. cloacae* induced an inflammatory response in monocytes, which was less present for *F. prausnitzii* and *B. ovatus* (**Figure S5G**). The *E. cloacae*-induced increase in IL-6 secretion could be reduced by pharmacological TLR5 inhibition (**Figure S5H**). These results collectively indicate that islet-resident macrophages play a major role in *E. cloacae*-induced islet-cell inflammation and dysfunction.

**Figure 4.**
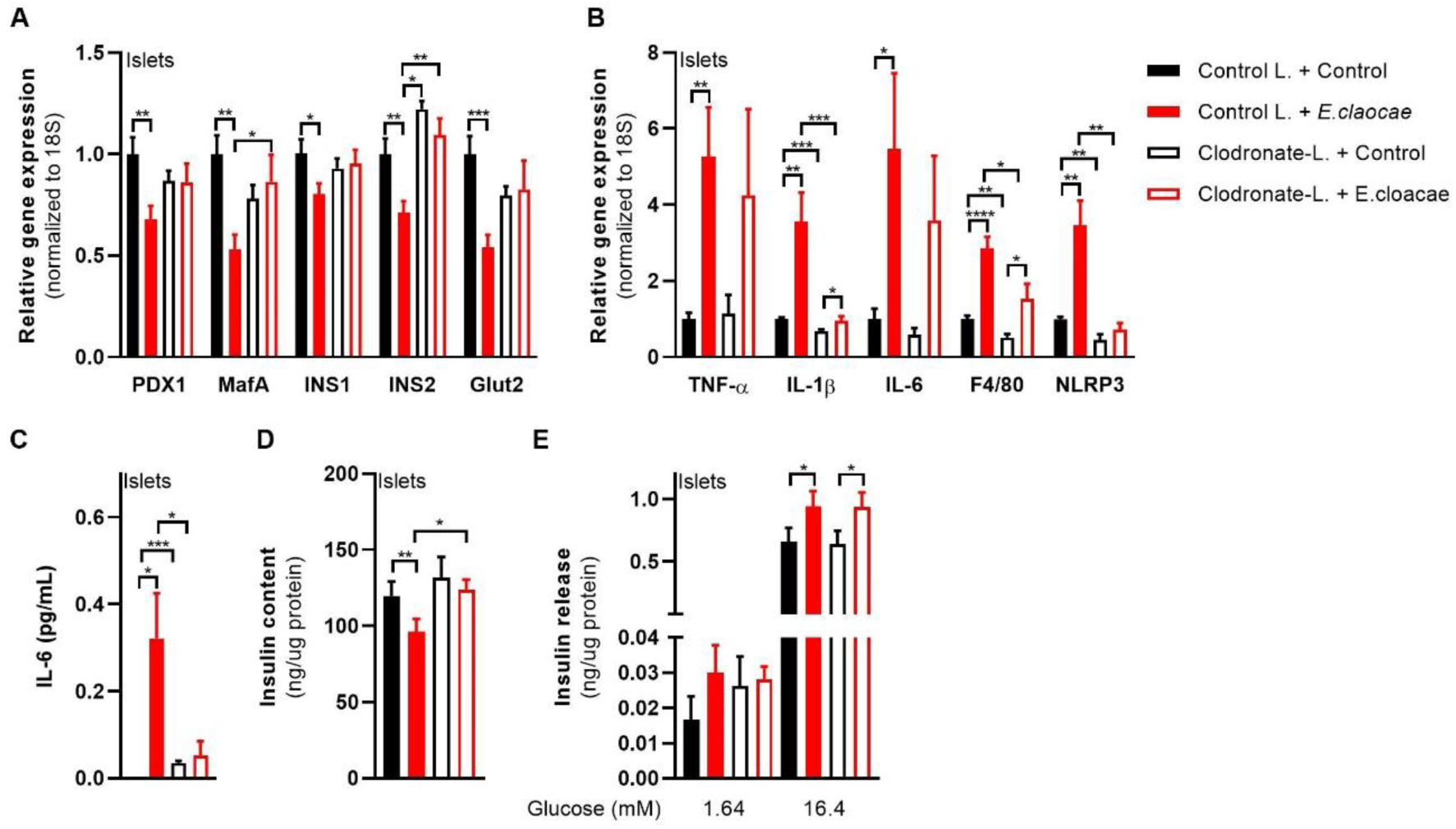
Macrophages mediate beta cell dysfunction in pancreatic islets. Macrophage-depleted islets from wild-type C57BL6J mice were treated with heat-inactivated *Enterobacter cloacae* (1E6 CFUs/mL) for 72h. (A) *E. cloacae* reduces beta-cell gene expression. Macrophage depletion in pancreatic islets protects from *E. Cloacae-*induced reduction in beta-cell gene expression. (B) *E. cloacae* increases beta-cell inflammation. Macrophage depletion in pancreatic islets reduces *E. Cloacae* induced inflammation. (C) *E. cloacae* increases beta-cell IL-6 secretion which is prevented by macrophage depletion in pancreatic islets. (D) *E. cloacae* reduces insulin content, which is prevented by macrophage depletion. (E) *E. cloacae* induces insulin hypersecretion both in WT and in macrophage depleted islets. Data shown are mean ± SEM. Unpaired t-test was used for statistical analysis (A-K). Significance level: *p<0.05, **p<0.01, ***p<0.001, ****p<0.0001 (mean ± SEM, 3 representative experiments per panel). Abbreviations: PDX1, pancreatic and duodenal homeobox 1; INS1 and INS2, insulin 1 and 2; NLRP3, NACHT, LRR and PYD domains-containing protein 3; TNF-α, tumor necrosis factor-alpha; IL-1β, Interleukin 1 beta; IL-6, Interleukin 6; TLR2, Toll-like receptor 2; TLR5, Toll-like receptor 5.

### Bacterial flagellin induces beta-cell inflammation and dysfunction

Flagellin is the main ligand for TLR5 (Hug et al., 2018) and therefore we hypothesized that this bacterial component could be the driving force behind the *E. cloacae*-induced phenotype. Indeed, both flagellin and flagellin-bearing *E. cloacae* activated TLR5 in a

human embryonic kidney (HEK) reporter cell line (**Figure 5A**), which was dose-dependently inhibited by the TLR5 inhibitor TH1020. Furthermore, flagellin induced a pro-inflammatory response in human macrophages, which was reduced when co-incubated with TH1020 (**Figure 5B**). Additionally, flagellin impaired insulin gene expression (**Figure 5C**), induced beta-cell inflammation (**Figure 5D-E**) and reduced insulin content (**Figure 5F**) while promoting insulin hypersecretion in pancreatic islets (**Figure 5G**), thus resembling the phenotype induced by *E. cloacae*.

**Figure 5.**
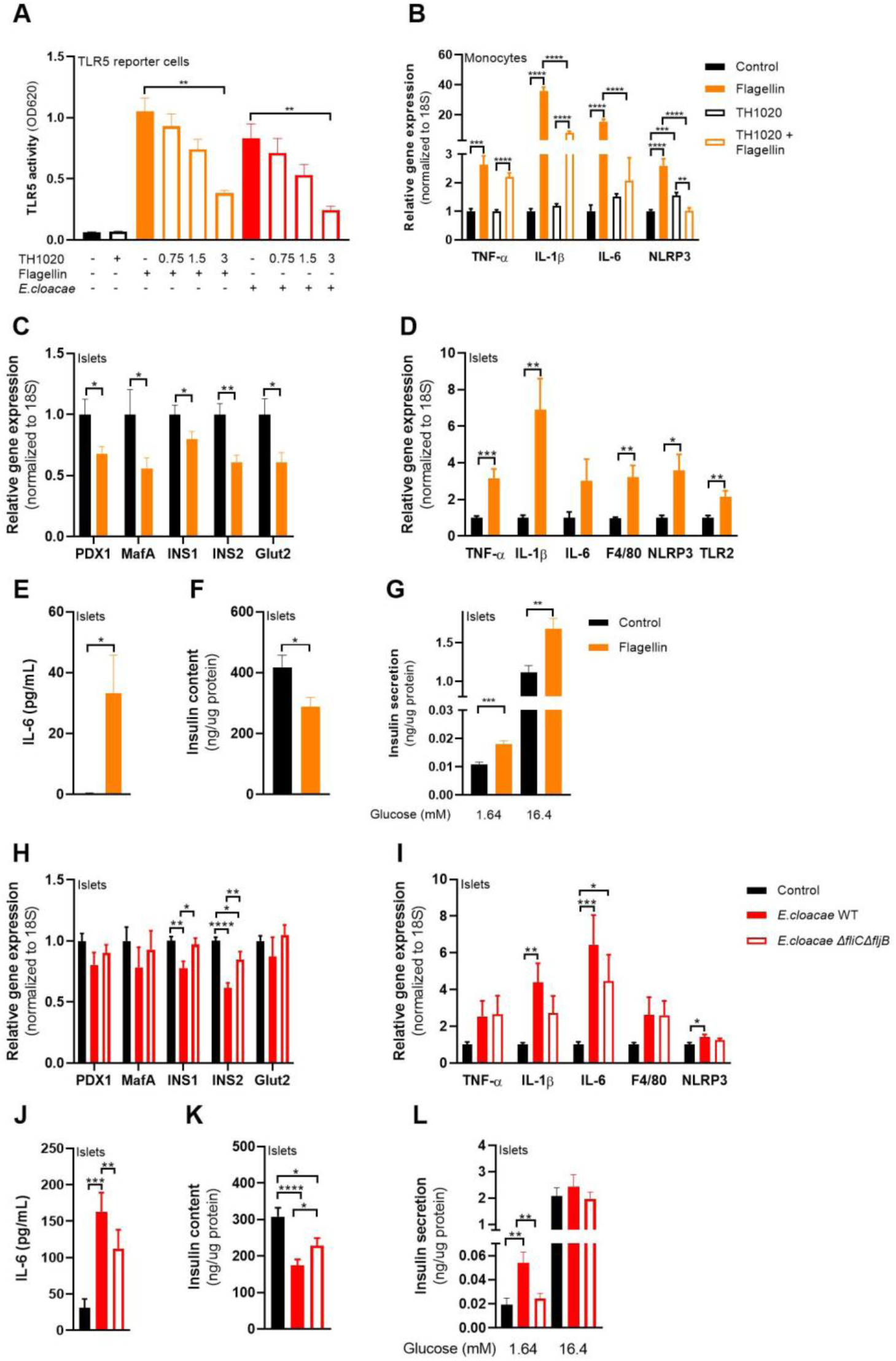
Flagellin induces beta cell dysfunction in pancreatic islets. HEK TLR5 reporter cell line (A), human monocytes (B) and pancreatic islets (C-L) from C57BL6J mice were incubated with flagellin (100 ng/mL), *E.cloacae* (1E6 CFUs/mL) or flagellin knock out *E.cloacae* (H-L). (A) Flagellin and *E. cloacae* activate TLR5 in HEK reporter cell line. TLR5 inhibitor TH1020 inhibits receptor activity dose-dependently (in µM). (B) Flagellin increases expression of inflammatory genes in macrophages, which is reduced by TLR5 inhibitor TH1020 (3 µM). (C) Flagellin reduces beta-cell gene expression. (D) Flagellin increases expression of inflammatory genes in islets. (E) Flagellin increases secreted IL-6 by islets. (F) Flagellin reduces insulin content in islets. (G) Flagellin induces insulin hypersecretion in islets. (H) *E. cloacae* reduces beta-cell gene expression. Knock out of flagellin in *E. Cloacae* partially protects against this loss of gene expression. (I) *E. cloacae* induces beta-cell gene inflammation. Knock out of flagellin in *E. Cloacae* partially protects against this inflammatory response. (J) *E. cloacae* stimulates IL-6 secretion by islets. Knock out of flagellin in *E. Cloacae* partially protects against this enhanced IL-6 release. (K) *E. cloacae* reduces beta-cell insulin content. Knock out of flagellin in *E. Cloacae* protects against this loss of insulin stores. (L) *E. cloacae* induces insulin hypersecretion at low glucose. Knock out of flagellin in *E. Cloacae* protects against this impaired secretory response. Data shown are mean ± SEM. Unpaired t-test (A-E, H-J) or Mann-Whitney (F, G, K, L) test was used for statistical analysis: *p<0.05, **p<0.01, ***p<0.001, ****p<0.0001. Abbreviations: INS1 and INS2, insulin 1 and 2; NLRP3, NACHT, LRR and PYD domains-containing protein 3; IL-1β, Interleukin 1 beta; IL-6, Interleukin 6; TLR, toll like receptor; ΔfliCΔfljB, flagelline genes knock out.

To further dissect the role of flagellin in the beta-cell deteriorating effects mediated by *E. cloacae*, we generated an *E. cloacae*-flagellin strain (Δ*fliC*Δ*fljB)* lacking both *fliC* and *fljB.* FliC and FljB are two different flagellar filament proteins, and homologous to those expressed by *Salmonella enterica* (Bonifield and Hughes, 2003). Compared to wild-type *E. cloacae*, *E. cloacae* Δ*fliC*Δ*fljB* did not suppress beta-cell gene transcription (**Figure 5H**). In addition, expression of inflammatory cytokines and secreted IL-6 were lower in islets exposed to the Δ*fliC*Δ*fljB* strain as compared to islets exposed to wild-type *E. cloacae* (**Figure 5I-J**). Islets had higher insulin content after incubation with the Δ*fliC*Δ*fljB* strain as compared to wild-type *E. cloacae* (**Figure 5K**). Finally, the Δ*fliC*Δ*fljB* strain did not induce fasting insulin hypersecretion (**Figure 5L**). Collectively, these results strongly suggest that flagellin, as part of the flagellum carried by bacteria belonging to Enterobacteriaceae, plays a pivotal role in beta-cell inflammation and beta-cell dysfunction via TLR5 activation on resident islet macrophages.

### Flagellin treatment augments insulin secretion in mice

To translate beta-cell dysfunction inducing effects of flagellin to an *in vivo* situation, we injected flagellin intraperitoneally into diet-induced obese (DIO) C57BL6J mice twice weekly for four weeks (**Figure 6A**). Flagellin injection did not alter body weight or fasting glucose (**Figure 6B-C**). However, flagellin-treated mice had lower glucose levels during an intraperitoneal glucose tolerance test compared to the placebo group (**Figure 6D-E**), which was driven by increased insulin secretion (**Figure 6F**) since insulin sensitivity did not differ between groups (**Figure 6G-H**). Similar to the *in vitro* experiments, there was a higher inflammatory tone in the pancreas of flagellin-treated mice as shown by higher *IL-1β* expression (**Figure 6I**) and a trend towards more inflammation in pancreatic islets isolated from flagellin-treated mice (**Figure 6K**). Insulin content of isolated islets did not differ between groups, while insulin release tended to increase during glucose-stimulated insulin secretion *ex vivo* in islets isolated form the flagellin group (**Figure 6J-M**). These results suggest that flagellin-induced transcriptional and functional alterations in beta-cell function, as observed *in vitro*, could be largely replicated *in vivo* in a mouse model.

**Figure 6.**
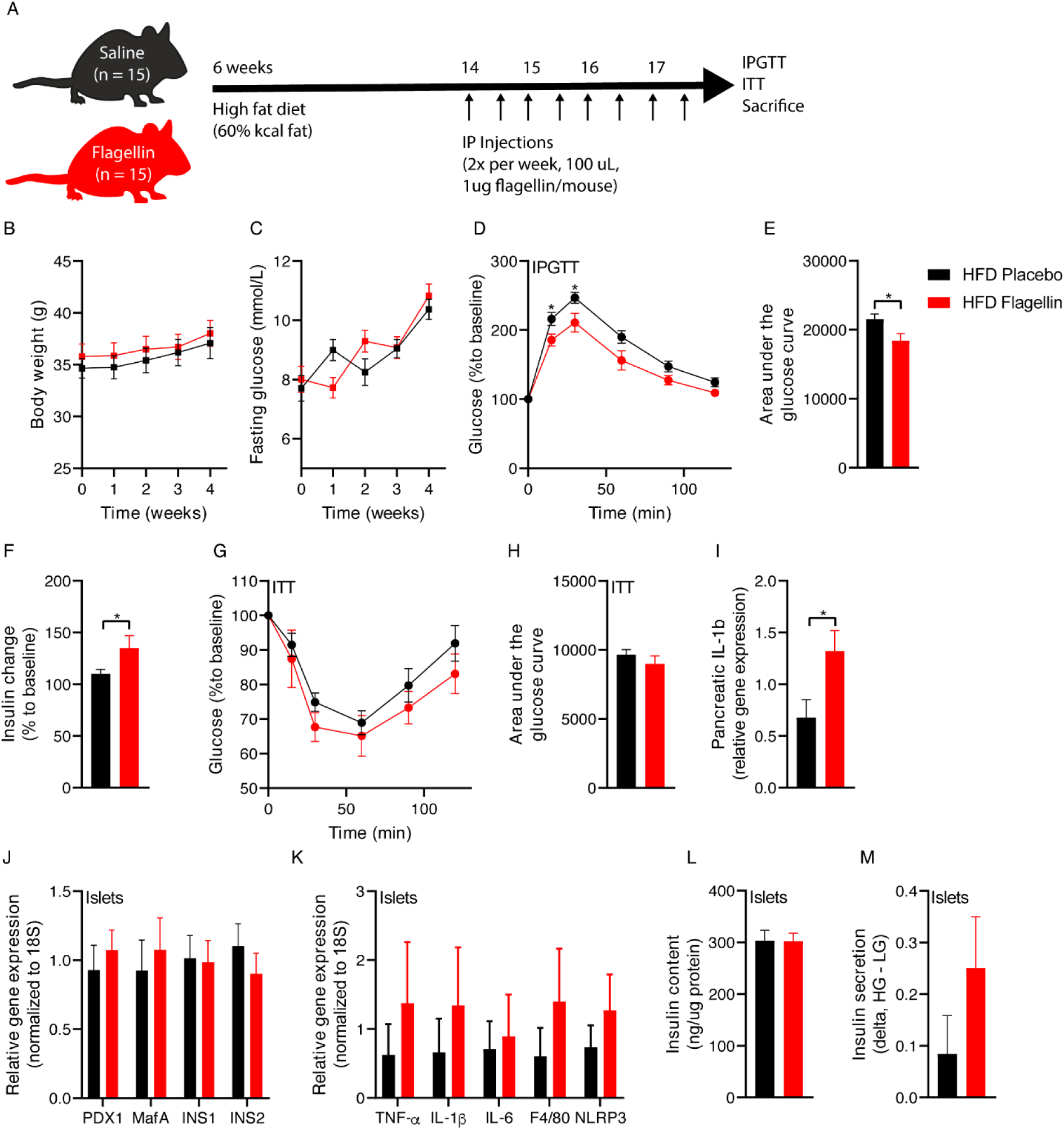
Flagellin injection in mice disturbs glucose tolerance. (A) Six-week old mice were fed a high fat diet (60%kcal fat) for 12 weeks. In the last 4 weeks of the diet, the mice were injected with either 1 ug flagellin in 100 uL saline or saline alone twice weekly. (B) Flagellin injections do not change the body weight. (C) Flagellin injections do not change fasting plasma glucose concentrations. (D, E) Flagellin-injected mice have improved glucose tolerance compared to placebo-treated mice (lower area under the glucose curve). (F) Fold change (15 min to baseline) of plasma insulin is greater in flagellin-treated mice compared to placebo (n = 10). (G, H) Insulin sensitivity is not affected by flagellin injection (0.75 IU/kg). The relative change in glucose to baseline is shown. (I) Flagellin increases pancreatic IL-1b expression. (J) Pancreatic islets were isolated from flagellin or saline treated mice, rested for 3 hours and gene expression was measured. Flagellin injections do not affect beta-cell gene expression (n = 5). (K) Flagellin injection numerically increases markers of beta cell inflammation (n = 5). (L) Insulin content was measured after islets were treated first with low glucose, followed by high glucose for 1h each. Flagellin injection does not affect insulin content (n =5) (M) Glucose stimulated insulin secretion was performed on islets and insulin release was measured at low as well as high glucose condition for 1h each. Flagellin injection numerically increases insulin release from beta cells (n = 5) Data shown are mean ± SEM. Unpaired t-test (E, F, I) or Šídák’s multiple comparisons test (D) was used *p<0.05, **p<0.01, ***p<0.001, ****p<0.0001. Abbreviations: INS1 and INS2, insulin 1 and 2; NLRP3, NACHT, LRR and PYD domains-containing protein 3; IL-1β, Interleukin 1 beta; IL-6, Interleukin 6; TNF-α, tumor necrosis factor-alpha; ITT, insulin tolerance test, IP, intraperitoneal; IPGTT, IP glucose tolerance test; HG, high glucose; LG, low glucose.

### Systemic flagellin dissemination relates to beta-cell dysfunction in humans

Fecal flagellin has been reported to be increased in obese people compared to lean controls (Tran et al., 2019). Similarly, we observed that obese mice had a higher flagellin load compared to lean mice (**Figure S6**). We predicted fecal flagellin gene abundance in the 150 HELIUS participants by inference from 16S rRNA profiles using PICRUSt (**Table S3**, **Figure 7A**). Fecal flagellin gene abundance was increased in T2D (**Figure 7B**). Next, we measured bacterial flagellin in the human blood circulation of the HELIUS cohort. While there was no difference between the matched obese groups, we did observe a positive correlation between serum flagellin load and HbA1c in T2D (**Figure 7C**). We hypothesized that increased flagellin reaches the circulation following a meal, which was previously shown to drive translocation of endotoxins (Ghoshal et al., 2009). We therefore measured postprandial plasma flagellin, C-peptide and plasma glucose in 80 matched participants of our bariatric surgery cohort (Van Olden et al., 2021) comprising obese normoglycemic and obese T2D people during a mixed-meal test (MMT) (**Figure 7A**, **Table S4**). C-peptide was chosen as it reflects insulin secretion rates and is not affected by potential differences in clearance by the liver, as is the case for plasma insulin concentrations. The MMT additionally allowed us to study the relationship between meal-induced flagellin and beta-cell response to an MMT. Participants with T2D had hyperglycemia following the MMT (**Figure 7D-E**) and lower C-peptide concentrations (**Figure 7F-G**) compared to normoglycemic humans with obesity. In both groups, flagellin increased during the MMT (**Figure 7H**). Postprandial area under the curve (AUC) for flagellin correlated with AUC C-peptide (**Figure 7I**), highlighting the link between bacterial flagellin and beta-cell insulin secretion in human beta-cell physiology.

**Figure 7.**
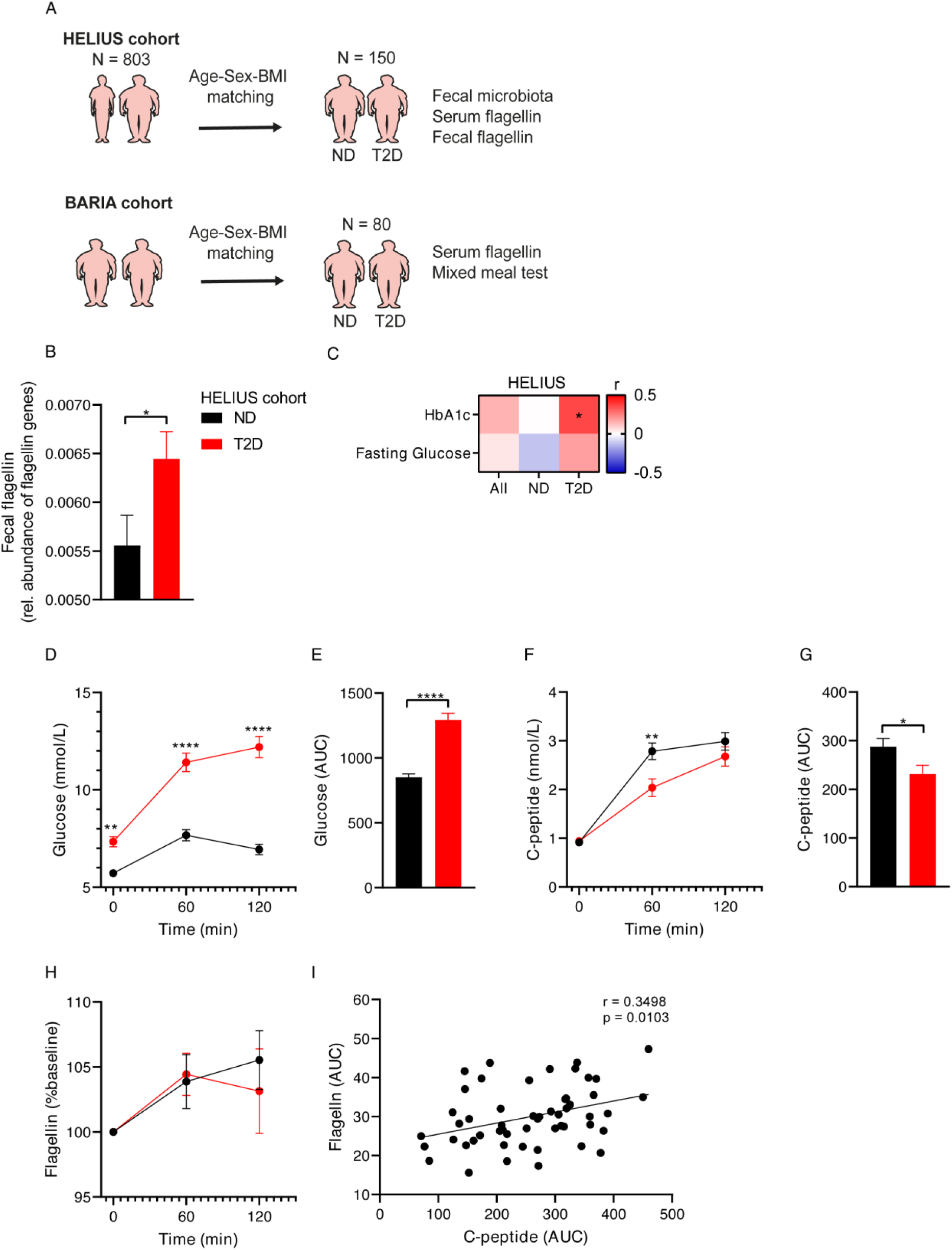
Fecal and serum flagellin is associated with glucose intolerance in humans. (A) 150 people were randomly selected from the HELIUS cohort (Snijder et al., 2017). Participants with T2D were matched with normoglycemic controls according to age, sex and BMI. In addition, 80 participants were selected from our bariatric surgery cohort (Van Olden et al., 2021). (B) Fecal flagellin genes are increased in T2D, as inferred from 16S rRNA gene profiles (HELIUS). (C) Serum flagellin positively correlates with HbA1c in people with T2D (HELIUS). (D, E) Plasma glucose concentrations during a mixed meal test (BARIA). Glucose levels are higher in people with T2D compared to normoglycemic obese controls. (F, G) Plasma C-peptide concentrations during a mixed meal test (MMT) (BARIA). C-peptide levels are higher in the obese normoglycemic controls versus T2D participants. (H) Serum flagellin increases during a mixed meal test (BARIA). (I) A positive correlation between serum flagellin area under the curve (AUC) and plasma C-peptide AUC during the MMT (BARIA) exists. Data shown are mean ± SEM. Unpaired t-test (B, E, G), Spearman correlation (C, I) and Šídák’s multiple comparisons test (D, F) was used: *p<0.05, **p<0.01, ***p<0.001, ****p<0.0001. Abbreviations: HELIUS, Healthy Life in an Urban Setting; AUC, Area under the curve.

## Discussion

In this study, we reveal a novel pathway by which flagellin, a structural component of notably Gram-negative bacteria residing in the gut, systemically disseminates following food ingestion. In pancreatic islets that are abundantly vascularized (Brissova and Powers, 2008), we propose that flagellin activates the innate immune system and induces an inflammatory response following binding to TLR5 receptors expressed by resident islet macrophages. This leads to beta-cell dysfunction, characterized by impaired insulin gene expression, impaired insulin processing, insulin hypersecretion/hyperinsulinemia and reduced insulin cell content. This study provides a new insight into the link between gut microbiota composition and T2D.

Beta-cell dysfunction is the key abnormality that leads to the development of hyperglycemia and T2D. Beta-cell dysfunction is characterized by inappropriate fasted or postprandial insulin secretion, which can either be excessive or insufficient, upon exposure to glucose or other nutrients (Johnson, 2021). While in the later stages of the disease insulin secretion rates are hampered, in people with prediabetes or early after diagnosis of T2D, hyperinsulinemia is often observed (Weyer et al., 2001). The role of hyperinsulinemia in T2D development has received ample attention. While initially reported to be a response against obesity-related insulin resistance, research has indicated that increased insulin secretion can develop in the absence of insulin resistance implying primary beta-cell pathology (Staimez et al., 2013). Importantly, hyperinsulinemia *per se* has negative effects.

Induction of hyperinsulinemia has been shown to promote obesity (Mehran et al., 2012), while prevention of hyperinsulinemia by pancreas-specific genetic knock out of insulin expression prevented obesity, improved insulin sensitivity and did not result in overt hyperglycemia (Mehran et al., 2012; Templeman et al., 2015; Templeman et al., 2017). With respect to the pancreatic islets, a chronic demand on beta cells to produce insulin is detrimental. As such, a prolonged increase in insulin secretory rates have been related to endoplasmic reticulum (ER) stress, depletion of intracellular insulin stores and beta-cell apoptosis (Hasnain et al., 2014). Pancreatic islets from individuals with T2D have lower insulin content compared to healthy controls (Cantley and Ashcroft, 2015; Henquin, 2019; Rahier et al., 2008; Rosengren et al., 2012). In mice, hyperglycemia leads to insulin content loss (Brereton et al., 2014). Reversibly, strategies that induce beta-cell rest are linked to improved beta-cell function over time (van Raalte and Verchere, 2017).

Current evidence relates overnutrition of carbohydrates and non-esterified fatty acids to beta-cell dysfunction (Esser et al., 2020). Another factor concerns a low-grade inflammatory response (Hajmrle et al., 2016). Beta-cell inflammation is a known hallmark of islets in people with T2D (Boni-Schnetzler and Meier, 2019) and a central role in this regard has been proposed for islet macrophages (Ehses et al., 2007). Macrophages are essential for normal beta-cell function and physiology (Nackiewicz et al., 2020), however, macrophages with a pro-inflammatory phenotype have been linked to beta-cell dysfunction (Nackiewicz et al., 2014). Triggers for activation of pro-inflammatory macrophages are uncertain but may involve hyperglycemia (Maedler et al., 2001), dyslipidemia (Igoillo-Esteve et al., 2010) and human islet amyloid polypeptide (hIAPP) (Westwell-Roper et al., 2014).

Here, we show an infectious stimulus triggering inflammation and beta-cell dysfunction: flagellin derived from intestinal microbiota. While Gram-negative bacteria are known to produce the canonical flagellin which is studied here, several other intestinal Firmicutes species are motile and have been described to contain flagella, including several *Roseburia*, *Clostridium*, and *Lactobacillus* spp (Dehoux et al., 2016; MM et al., 2015; Tamanai-Shacoori et al., 2017). However, the flagellins of these latter, Gram-positive bacteria have not been well characterized and some are glycosylated resulting in attenuated TLR5 signaling efficiency (Kajikawa et al., 2016).

A first link towards flagellin came from the observation that in a large cohort the fecal abundance of the family of Enterobacteriaceae, specifically *E. cloacae*, was increased in people with T2D and that the fecal abundance of Enterobacteriaceae and *E. cloacae* correlated with glucose intolerance in humans. Administration of *E. cloacae* by oral gavage to mice fed a high-fat diet has previously been shown to induce glucose intolerance (Fei and Zhao, 2013). A proposed mechanism by which gut microbiota may influence host metabolism is by escaping immune control and translocating to extra-intestinal tissues (Amar et al., 2011a; Amar et al., 2011b). While this has been shown for adipose tissue (Massier et al., 2020; Udayappan et al., 2017), translocation of intestinal bacteria into the pancreas was also suggested to trigger the influx of immune cells and islet inflammation (Thomas and Jobin, 2020). In patients undergoing pancreatoduodenectomy, pancreatic fluid contained bacterial DNA, with a similar composition, density and diversity as bile and jejunal fluid (Rogers et al., 2017), suggesting direct translocation from the small intestine into pancreatic juice. Others also suggest a bacteriome (Riquelme et al., 2019) and mycobiome (Aykut et al., 2019) in pancreatic tissue of cancer patients. In line with these data, we observe a correlation between systemic antibodies against *E. cloacae* and hyperglycemia, which may suggest translocation of at least parts of this bacteria to extraintestinal sites. Nevertheless, translocation of whole bacteria remains rather controversial (Scheithauer et al., 2020) since there are major challenges related to the sequencing of small amounts of bacterial DNA in extraintestinal tissues (de Goffau et al., 2019).

Heat-inactivated *E. cloacae* induced a detrimental beta-cell phenotype with insulin hypersecretion, induction of ER stress markers (elevated PI/I ratio), inflammation and reduced beta-cell insulin content. We show that deletion of TLR5, of which flagellin is the dominant ligand, on resident islet macrophages protected against the effects of the flagellum-bearing *E. cloacae*. Flagellum is a virulence factor that enables bacteria to move within the intestine and even adhere to the intestinal wall, a process called encroachment (Haiko and Westerlund-Wikström, 2013; Tran et al., 2019). Flagellin fully reproduced the beta-cell phenotype of *E. cloacae*. A causal role for flagellin was observed in *E. cloacae* with flagellin knock out, where the effects on inflammation, hypersecretion and reduced insulin content were strongly diminished. Strengthening the role of flagellin, mice that were injected with flagellin exhibited a similar beta-cell phenotype.

In line with our previous work (Scheithauer et al., 2021), we observed an increment in plasma flagellin concentrations following a meal in the current cohort. This indicates that flagellin, like the widely studied LPS, may translocate after food ingestion. Importantly, flagellin is able to pass the epithelial barrier (Gewirtz et al., 2001) and is a potent stimulus of the mucosal immune response (Cullender et al., 2013a; Vijay-Kumar and Gewirtz, 2009). Plasma flagellin levels also positively correlated with HbA1c in our cohort, while the meal-related flagellin increment associated with higher C-peptide release in obese humans with or without diabetes.

Finally, to support the hypothesis that systemically disseminated flagellin causes the observed beta-cell phenotype, we collected pancreatic biopsies from people with T2D undergoing pancreatic surgery for benign lesion. We collected five biopsies of the head of the pancreas. Recent antibiotics use (<3 months) was an exclusion criterion (**Table S5**). While it was technically not possible to measure flagellin in these biopsies due to interference of the pancreatic enzymes with the flagellin assay, we observe the presence of antibodies against flagellin, supporting the notion that the well-vascularized pancreatic islets are exposed to flagellin (**Figure S7**). A previous study indicated that a functional immune response is essential to control flagellin expression bacteria (Cullender et al., 2013a), which seems to be reduced in obese humans (Tran et al., 2019).

We acknowledge a number of limitations of this study. First, although the concept of translocation of flagellin to the systemic circulation seems to be plausible to explain beta-cell inflammation (Thomas and Jobin, 2020), we can only speculate if bacterial flagellin is transported via the blood circulation towards the pancreas. More sensitive methods are necessary to quantify small amounts of bacterial components such as flagellin in extra-intestinal tissues (Scheithauer et al., 2020; Tran et al., 2019). Second, we provide evidence that people with T2Ds have a higher fecal and circulating flagellin load compared to normoglycemic individuals, with flagellin loads correlating with hyperglycemia. However, such a correlation does not show causation. Future research should evaluate whether reducing intestinal or systemic flagellin load will improve beta-cell function and can reduce diabetes incidence. Third, the concept of insulin hypersecretion and its linked to low-grade inflammation needs further validation; as such, studies that show the benefits of reducing insulin hypersecretion are currently scant. In addition, a moderate inflammatory response has been suggested to be beneficial in stimulating insulin release (Ying et al., 2020), although the effects of prolonged inflammation may have different effects.

Together, we present a novel pathway linking bacterial flagellin from the Gram-negative *E. cloacae* in a TLR5-macrophage-dependent manner to beta-cell inflammation and beta-cell dysfunction, suggesting a new mechanism linking gut microbiota and T2D prevalence and opening up potential avenues for novel therapies.

## Acknowledgments

This study was funded by Diabetes Fonds (Application number: 2015.81.) and Marie Skłodowska-Curie Actions (Call H2020-MSCA-IF-2015). CBV was supported by CIHR project grant PJT-165943. M.N. is supported by a personal ZONMW VICI grant 2020 [09150182010020]. B.A.V. is the Children with Intestinal and Liver Disorders (CH.I.L.D) Foundation Chair in Pediatric Gastroenterology. The HELIUS study is conducted by the Amsterdam University Medical Centers, location AMC and the Public Health Service of Amsterdam. Both organisations provided core support for HELIUS. The HELIUS study is also funded by the Dutch Heart Foundation, the Netherlands Organization for Health Research and Development (ZonMw), the European Union (FP-7), and the European Fund for the Integration of non-EU immigrants (EIF). We are most grateful to the participants of the HELIUS study and the management team, research nurses, interviewers, research assistants and other staff who have taken part in gathering the data of this study. The study reported here was additionally supported by (an) additional grant(s) from Dutch Heart Foundation: 2010T084 (K Stronks), ZonMw: 200500003 (K Stronks), European Union (FP-7): 278901 (K Stronks), European Fund for the Integration of non-EU immigrants (EIF): 2013EIF013 (K Stronks). HH is supported by a Senior Fellowship of the Dutch Diabetes Research Foundation (2019.82.004).

## Author contributions

T.P.M.S. performed the experiments and prepared the manuscript. M.W. performed experiments. S.R.H., G.S., M.S. and D.D. assisted with animal experiments. S.M., M.dB., A.vdL. and Ö.A. conducted the BARIA cohort. M.B. and M.D. performed bioinformatic analyses. W.M.dV., C.B., H.Y. and C.M. provided bacterial cultures. M.G.B and O.R.B. provided pancreatic biopsies. G.M.D.T., G.J.B. and W.M.V. aided with the writing. B.J.H.B. conducted the HELIUS cohort. B.A.V., M.N., H.H. and C.B.V. supervised the project. D.H.R. developed the theory and supervised the project.

## METHODS

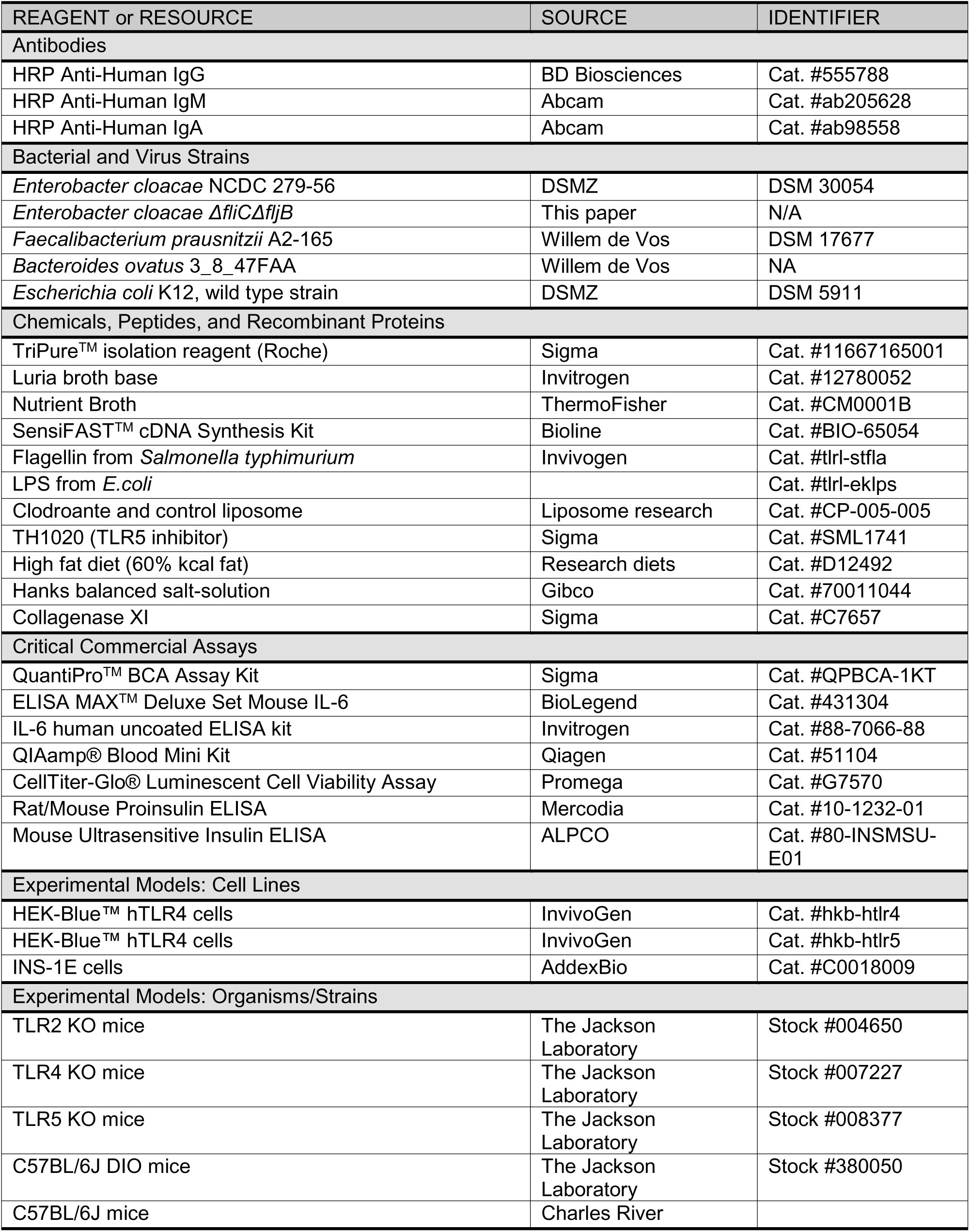

## LEAD CONTACT AND MATERIALS AVAILABILITY

Further information and requests for resources and reagents should be directed to and will be fulfilled by the Lead Contact, Daniel H. van Raalte or Torsten P.M. Scheithauer. For bacterial mutants, please contact Bruce A. Vallance

## EXPERIMENTAL MODEL AND SUBJECT DETAILS

### Participants

For the current study we included 803 people of Dutch descent with available data on the gut microbiome from the HELIUS cohort in Amsterdam the Netherlands (Snijder et al., 2017). For details, regarding the HELIUS study (recruitment, data collection) in general, and regarding this selection in particular, see Deschasaux et al. (2018). For microbiome analysis, 150 participants were randomly selected and diabetic participants (n = 100) were age-BMI-sex matched to healthy non-diabetic controls (n = 50). Diabetic participants were selected according to one of the following criteria (at least one): self-reported diagnosis of T2D, use of antidiabetic medication, fasting blood glucose > 7.0 mmol/L and HbA1c > 48 mmol/mol. All participants did not use antibiotics for the last 3 months.

The HELIUS data are owned by the Amsterdam UMC, location AMC in Amsterdam, The Netherlands. Any researcher can request the data by submitting a proposal to the HELIUS Executive Board as outlined at http://www.heliusstudy.nl/en/researchers/collaboration, by The HELIUS Executive Board will check proposals for compatibility with the general objectives, ethical approvals and informed consent forms of the HELIUS study. There are no other restrictions to obtaining the data and all data requests will be processed in the same manner.

To validate our results in the HELIUS cohort, we included 40 T2D participants and 40 non-diabetic, age, sex and BMI matched controls of the BARIA cohort. For details, see Van Olden et al. (2021). The BARIA Study aims to assess how microbiota and their metabolites affect transcription in key tissues and clinical outcome in obese subjects and how baseline anthropometric and metabolic characteristics determine weight loss and glucose homeostasis after bariatric surgery.

The studies were approved by the local Institutional Review Board of the Amsterdam UMC, location AMC in Amsterdam, the Netherlands, and conducted in accordance with the Declaration of Helsinki.

### Bacteria

*Enterobacter cloacae* NCDC 279-56 and *Escherichia coli* K12 were cultured in Luria broth base (Invitrogen, US) and on LB Agar (Invitrogen, US) at 37°C, overnight, before being used for experiments. *Faecalibacterium prausnitzii* A2-165 in YCFA media and *Bacteroides ovatus* 3_8_47FAA in YZFAA media.

### Animals

C57BL/6J mice were purchased from Charles River (France) and maintained under specific pathogen free conditions in the S-building of the Amsterdam UMC, location AMC. TLR2 KO, TLR4 KO, TLR5 KO and C57BL/6J DIO mice were purchased from Jackson Laboratory (JAX); control animals on C57BL/6J background were used from JAX facilities instead of Charles River. All animals were socially housed, under a 12h light/dark cycle until 12-14 weeks and sacrificed for pancreatic islets isolation. Only male mice were included in this study. Animal work was performed in accordance with the Central Commission for Animal Experiments (CCD, The Netherlands).

## METHOD DETAILS

### Reagents and Antibodies

Luria broth base (Invitrogen, US), LB Agar (Invitrogen, US), sterile PBS (Fresenius Kabi, Germany), pentobarbital (EUTANASIA), collagenase XI (Sigma-Aldrich, US), Hanks balanced salt-solution (HBSS w/o calcium and magnesium, Gibco, US), RPMI 1640 (Gibco^TM^, US), fetal bovine serum (FBS, Capricorn, Germany), Penicillin-Streptomycin (P/S, Gibco, US), bovine serum albumin (BSA, RIA grade, Sigma, US), RIPA lysis buffer (ThermoScientific^TM^, US), TriPure^TM^ isolation reagent (Roche, Switzerland), GlycoBlue^TM^ (Invitrogen^TM^, US), UltraPure^TM^ DNase/RNase-Free Distilled Water (Invitrogen^TM^, US), SensiFAST^TM^ cDNA Synthesis Kit (Bioline, UK), SensiFAST^TM^ SYBR® No-ROX Kit (Bioline, UK), DMEM (high glucose, Gibco^TM^, US), β-Mercaptoethanol (Sigma, US), sodium pyruvate (Gibco^TM^, US), HRP Anti-Human IgG (BD Biosciences, US), HRP Anti-Human IgM (Abcam, UK), HRP Anti-Human IgA (Abcam, UK), Tween20 (Merck), 1-Step^TM^ Ultra TMB-ELISA Substrate Solution (ThermoScientific^TM^, US), mouse ultrasensitive insulin ELISA (ALPCO, US), QuantiPro^TM^ BCA Assay Kit (Sigma, US), ELISA MAX^TM^ Deluxe Set Mouse IL-6 (BioLegend, US), IL-6 human uncoated ELISA kit (Invitrogen^TM^, US), clodronate and control liposome (Liposome research, The Netherlands), TH1020 (Sigma, US), CellTiter-Glo® Luminescent Cell Viability Assay (Promega, US), Rat/Mouse Proinsulin ELISA (Mercodia, SE).

### Heat-inactivation of bacteria

The optical density of the bacterial culture was measured at 600 nm (OD600) and diluted to 1E9 colony forming units (CFUs) per mL. Bacteria were centrifuged at 8000 xg for 5 minutes and resuspended in 1 mL sterile phosphate buffered saline (PBS). All bacteria were heat-inactivated at 70°C for 30 min and stored at -80°C in small aliquots for further use.

#### Pancreatic surgery

Individuals who are scheduled for pancreatic surgery (e.g., pylorus-preserving pancreatoduodenectomy or Whipple’s procedure), because of pancreatic carcinoma, were asked to donate healthy tissue surrounding the tumor. Tissue was harvested under surgical conditions, snap frozen in liquid nitrogen and stored at -80°C until further analysis.

### Pancreatic islet isolation

Mice were anaesthetized with 2.5 mg pentobarbital (diluted in sterile saline) per mouse and sacrificed via cervical dislocation. After clamping the *Ampulla of Vatar*, the pancreas was injected intraductally with approximately 3 mL of collagenase XI (1000 U/ml) in HBSS (without calcium chloride) and placed in 50 mL tubes with an additional 2 mL of collagenase solution. The pancreas was incubated at 37°C for 13 minutes followed by gentle shaking to obtain a homogenously dispersed pancreas. Digestion was stopped with cold HBSS supplemented with 1 mM CaCl_2_. Islets were washed two times in cold HBSS with CaCl_2_ by centrifuging 185 xg for 30 seconds. Next, islets were filtered through a 70 μM prewetted cell strainer. After flushing two times with 10 mL of HBSS with CaCl_2_, the strainer was turned upside-down over a Petri dish and rinsed with 16 mL of islet media (RPMI 1640 with GlutaMAX^TM^ 1x, 10% FBS and P/S 1x) to collect the islets into the dish. Islets were handpicked under the Nikon SMZ800 microscope into a fresh Petri dish with islet media. Islets were rested overnight to recover from isolation procedure.

### Plasmid construction

Overlap extension PCR (Ho et al., 1989) was used to generate pRE118-pheS-ΔfliC and pRE118-pheS-*ΔfljB* constructs (pRE118-pheS was a gift from Christopher Hayes of UC Santa Barbara). For pRE118-pheS-*ΔfliC* construct, two PCR fragments were amplified using *E. cloacae* genomic DNA as the template. Primer pairs used to amplify the PCR fragments are ecFliC-P1 (5’-GATGATGGTGATGGTACGCGTGGTACCGGTAGTCGCT-3’) plus ecFliC-P2 (5’-GGTTTCTAGGGTCGGTGCCTTAACACTCA-3’), and ecFliC-P3 (5’-CACCGACCCTAGAAACCCTGTCTCTGCTGCGTTAA-3’) plus ecFliC-P4 (5’-GACAGTGAGCTCGCATCGTTAACGCGTCTTCACCAA-3’), respectively. This results in a 789-bp fragment containing the upstream of *fliC* and a 750-bp fragment containing the downstream of the *fliC*, respectively. These two PCR fragments were then mixed and used as the template for a secondary PCR (with primer pairs ecFliC-P1 containing a KpnI restriction enzyme site and ecFliC-P4 containing a SacI restriction enzyme site). The 16-bp overlapping sequence (underlined) in primers ecFliC-P2 and ecFliC-P3 allows the amplification of a 1,539-bp PCR product. This PCR product was digested with KpnI and SacI, and directly cloned into the *E. cloacae* suicide vector pRE118-pheS (Kanr).

The pRE118-pheS-Δ*fliC* construct was generated the same as above. Primer pairs used to amplify the PCR fragments are FljB-P1 (5’-GCACGTCTAGAGTGACCTTTATCGTCATCTCACCGT-3’) plus FljB-P2 (5’-GTACCCAGCTGAGTCTGGGATTTGTTCAGGTTGTT-3’), and FljB-P3 (5’-AGACTCAGCTGGGTACTGCTGCGTTAATCTGCGTTA-3’) plus FljB-P4 (5’-GACAGTGAGCTCGTACAGCTATTCGCTGCATAACGA-3’), respectively. This results in a 955-bp fragment containing the upstream of *fljB* and a 950-bp fragment containing the downstream of the *fljB*, respectively. These two PCR fragments were then mixed and used as the template for a secondary PCR (with primer pairs FljB-P1 containing a XbaI restriction enzyme site and FljB-P4 containing a SacI restriction enzyme site). The 16-bp overlapping sequence (underlined) in primers FljB-P2 and FljB-P3 allows the amplification of a 1,905-bp PCR product. This PCR product was digested with XbaI and SacI, and directly cloned into the E. cloacae suicide vector pRE118-pheS.

### Generation of E. cloacae mutant strains

pRE118-pheS-*ΔfliC* and pRE118-pheS-*ΔfljB* constructs were transformed into *E. coli* MFD(λ pir). E. coli MFD(λ pir) carrying these constructs and WT *E. cloacae* were grown overnight in LB, and then mixed at a ratio of 4:1 (donor vs recipient strains). To make fljB fliC, E. coli MFD(λ pir) carrying pRE118-pheS-*ΔfljB* and *ΔfliC* were grown overnight in LB, and then mixed at a ratio of 4:1. Fifty microliter of the mixture was spotted onto LB agar plate containing diaminopimelic acid (DAP, 0.3 mM), and incubated at 37°C overnight. This was followed by scaping the cell mixtures in PBS and plating onto LB agar containing streptomycin (100 μg/ml) and kanamycin (50 μg/ml). The resulting single-crossover mutants were grown statically in LB at 37°C overnight, and further counter selected on M9 minimal medium agar plates containing 0.4% (w/v) glucose and 0.1% (w/v) p-chlor-ophenylalanine (Ting et al., 2020). Kanamycin sensitive colonies were screened by colony PCR. The ΔfliC deletion mutant was confirmed by PCR with primers ecFliC-check-F (5’-GCGTTTCTGATGGCGTTCTGAA-3’) and ecFliC-check-R (5’-GCTCGAACTTGTTCATCCCGATT-3’). The predicted size of WT and mutant bands is 1201-bp and 362-bp, respectively. The *ΔfljB* deletion mutant was confirmed by PCR with primers FljB-check-F (5’-GCAGAACAACCTGAACAAATCCCA-3’) and FljB-check-R (5’-GACACGTTTACGCCGGTTCACTAT-3’). The predicted size of WT and mutant bands is 1811-bp and 387-bp, respectively.

### Confirming mutants with a swimming motility assay

WT and mutant E. cloacae strains (*ΔfljB, ΔfliC, ΔfljBΔfliC*) were grown statically in 2 μl of LB at 30 °C for 18 h. Two microliter of these cultures were spotted onto semi-solid nutrient broth (BD) agar plates containing 0.3 % agar. After incubating the plates at 37°C for 4 h, pictures showing the swimming motility were taken (**Figure S8**).

### Glucose stimulated insulin secretion

Pancreatic islets or β-cell lines (see seeding below) were washed in a 12 well plate 2x with 500 uL low glucose Krebs-Ringer buffer (KRB; 132 mM NaCl, 5 mM KCl, 1 mM KH_2_PO_4_, 1 mM MgSO_4_, 2.5 mM CaCl_2_, 5 mM NaHCO_3_, 10 mM HEPES, 0.25% BSA, 1.64 mM glucose) and starved in 500 uL low glucose KRB for 1 hour. Islets were split into 10 islets per well in a 12 well plate (triplicates) and incubated for 1 hours in 500 uL low glucose KRB. The same islets were transferred into 500 uL high glucose KRB (16.4 mM) for 1 hour. Finally, islets were washed 2x with 1 mL PBS and lysed with 150 uL RIPA buffer. Islet lysate was spun at 14.000 xg for 10 min at 4°C and the supernatant was stored at -20°C until further use.

### DNA isolation

Fecal DNA was extracted from 150 mg fecal material and the sorted fractions using a repeated bead beating protocol (method 5) (Costea et al., 2017). DNA was purified using Maxwell RSC Whole Blood DNA Kit. 16S rRNA gene amplicons were generated as described below.

### RNA isolation and cDNA synthesis

RNA was isolated with TriPure^TM^ isolation reagent (Roche). Cells were separated from the culture media and 300 uL TriPure^TM^ was added. After lysis, 60 uL chloroform was added, the mixture was vigorously shaken for 15 seconds and incubated for 3 minutes at room temperature. Next, samples were spun for 15 minutes at 12.000 xg (4°C) and the aqueous phase was mixed with 190 uL isopropanol with 0.44 uL GlycoBlue^TM^. After an overnight incubation at -20°C, samples were spun at 12.000 xg for 10 minutes (4°C) and the pellet was 2x washed with 1 mL of 75% ethanol (7.500 xg, 5 minutes, 4°C). Next, the pellet was dried at room temperature for 10 minutes, 18 uL RNase free H2O was added and incubated at 56°C for 10 minutes. RNA concentration was measured with Nanodrop. cDNA synthesizes was performed with SensiFAST^TM^ cDNA Synthesis Kit according to manufactures instructions.

### PCRs

Gene expression was measured *via* real time quantitative PCR (RT-qPCR) with the aid of PCR machine (BioRad, US). SensiFAST^TM^ SYBR® No-ROX Kit was used according to manufactures instructions. For each well, 7.5 ng cDNA and 1 μM primer mix were used in a 10 uL PCR mix. For primers see **Table S6**. Temperatures are used as following, if not stated differently: 95°C for 10 minutes, 40 cycles of 95°C for 15 seconds and 60°C for 30 seconds with a plate reading, followed by a melt curve with increment of 0.5°C every 5 seconds starting from 65°C to 95°C.

Fecal bacteria were measured via quantitative PCR with the aid of PCR machine (BioRad, US). SensiFAST^TM^ SYBR® No-ROX Kit was used according to manufactures instructions. For each well, 10 ng genomic DNA and 300 nM primer mix were used in a 10 uL PCR mix. For primers see **Table S6**. For total bacterial in feces, EUBAC primers and temperature settings were used as stated (Nadkarni et al., 2002). For Enterobacteriaceae detection, En-lsu3 was used as described (Matsuda et al., 2007). Primers for *Enterobacter cloacae* was designed for the V3V4 regions. Temperatures as described above were used. Standard amplicons were made with genomic DNA from *E.coli* or *E. cloacae* and *Taq* DNA Polymerase (Qiagen, Germany) according to manufactures instructions. Amplicons were cleaned with QIAquick PCR purification Kit (Qiagen, Germany). Copy numbers were calculated according to the standard curve.

### Cell lines

HEK-Blue^TM^ hTLR5 cells were used according to manufactures instructions. For flagellin detection in the blood circulation, 20 uL serum was used per well (96 well plate) and mixed with 180 uL of 1.4×1E5 cells/mL in detection media. Cells were incubated for 16 hours and the supernatant was read at OD620. INS-1E cells were cultured in RPMI 1640 media (5% FBS, 1x P/S, 1x HEPES, 50 μM β-mercaptoethenol, 1x sodium pyruvate) and passaged with 0.25% Trypsin-EDTA. Cells were seeded in a 12 well plate at 75.000 cells/mL, rested overnight and incubated with heat-inactivated bacteria for 72 hours.

### Antibody analysis

Bacteria were grown overnight and the optical density was measured at OD600. Bacteria were diluted to have 1E9 CFUs/mL and washed with 1 mL sterile PBS (8000 xg, 5 minutes, 4°C). Bacteria were sonicated on ice at 30% amplitude for 20 x 30 seconds cycles with 60 seconds intervals. Nunc^TM^ MicroWell^TM^ 96-well microtiterplates (ThermoScientific^TM^, US) were coated with 200.000 sonicated bacteria per well (100 uL) overnight at 4°C. Plates were washed 3x with 300 uL per well of PBS and blocked with 150 uL PBS with 1% BSA for 2 hours at room temperature. Plates were washed again with PBS, 100 uL of 250x diluted serum samples (PBS/BSA) was added and incubated for 4 hours at room temperature. Plates were washed 3x with PBS with 0.05% Tween20 and 100 uL of secondary antibody (2000x diluted HRP anti-Human IgG; 50.000x diluted HRP anti-human IgM; 20.000x diluted HRP anti-human IgA; in PBS/Tween20) was added for 2 hours at room temperature. Plates were washed with PBS again and 100 uL of TMB was added for 15 minutes. The reaction was stopped with 50 uL of 0.5M HCl and read at OD450.

Pancreatic biopsies were 10x diluted according to tissue weight and homogenized in ultrapure water (Invitrogen, US) with the aid of a sterile metal bead. The homogenate was spun Nunc^TM^ MicroWell^TM^ 96-well microtiterplates (ThermoScientific^TM^, US) were coated overnight (4°C) with 100 ng per well of flagellin from *Salmonella typhimurium* (Invivogen, US). The plates was washed 3x with 300 uL PBS/Tween20. Afterwards, 100 uL of homogenized pancreas was added and incubated for 1h at 37°C. The plates was washed 3x with 300 uL PBS/Tween20. Secondary antibodies and TMB were added as described above.

### ELISA

Insulin was measured in low glucose KRB, high glucose KRB and cell lysate from GSIS experiments with ALPCO mouse ultrasensitive insulin ELISA according to manufactures instructions. Concentrations were normalized to total protein content measured via QuantiPro^TM^ BCA Assay Kit. Proinsulin ELISA (Mercodia) was performed according to manufactures instructions. IL-6 concentrations were measured in cell supernatants *via* ELISA MAX^TM^ Deluxe Set Mouse IL-6 and IL-6 human uncoated ELISA kit according to manufactures instructions.

### Monocyte isolation

PBMCs were isolated with Lymphoprep (GE Healthcare) and CD14 MACS beads (Miltenyi) according to manufactures instructions.

### Macrophage depletion

Pancreatic islet macrophages were depleted with Clodronate-liposome. Islets were isolated and rested for 3 hours. Islets were picked in a small petri dish (40-70 islets per dish) and treated with either clodronate or control liposome for 48h (1 in 5 diluted in

islet media). Islets were washed 3x with 2 mL complete media and picked in fresh media.

### Library preparation and sequencing

Library preparation and sequencing was performed at the Wallenberg Laboratory (Sahlgrenska University of Gothenburg, Sweden). Fecal microbiome composition was profiled by sequencing the V4 region of the 16S rRNA gene on an Illumina MiSeq instrument (Illumina RTA v1.17.28; MCS v2.5) with 515F and 806R primers designed for dual indexing (Kozich et al., 2013) and the V2 Illumina kit (2×250 bp paired-end reads). 16S rRNA genes from each sample were amplified in duplicate reactions in volumes of 25 µL containing 1x Five Prime Hot Master Mix (5 PRIME GmbH), 200 nM of each primer, 0.4 mg/ml BSA, 5% DMSO and 20 ng of genomic DNA. PCR was carried out under the following conditions: initial denaturation for 47 min at 94°C, followed by 25 cycles of denaturation for 45 sec at 94°C, annealing for 60 sec at 52°C and elongation for 90 sec at 72°C, and a final elongation step for 10 min at 72°C. Duplicates were combined, purified with the NucleoSpin Gel and PCR Clean-up kit (Macherey-Nagel) and quantified using the Quant-iT PicoGreen dsDNA kit (Invitrogen). Purified PCR products were diluted to 10 ng/μL and pooled in equal amounts. The pooled amplicons were purified again using Ampure magnetic purification beads (Agencourt) to remove short amplification products; for negative controls, see Deschasaux et al. (2018). Libraries for sequencing were prepared by mixing the pooled amplicons with PhiX control DNA purchased from Illumina. The input DNA had a concentration of 3 pM and contained 15% PhiX and resulted in the generation of about 700K clusters/mm2 and an overall percentage of bases with quality score higher than 30 (Q30) higher than 70%.

### Bioinformatic pipeline

USEARCH (v11.0.667_i86linux64) was used to process the raw sequencing reads. For paired-end merging, we used 30 max. allowed differences in the overlapping region (“maxdiffs”) for the merging step (using the “fastq_mergepairs” command) and max. 1 expected errors (“fastq_maxee”) as a quality filter threshold (using the “fastq_filter” command). Expected error-based read quality filtering is described in detail in Edgar et al. 2015. After merging paired-end reads and quality filtering, remaining contigs were dereplicated and unique sequenced were denoised using the UNOISE3 algorithm in order to obtain Amplicon Sequence Variants (ASVs). All merged reads were subsequently mapped against the resulting ASVs to produce an ASV table. ASVs not matching expected amplicon length were filtered out (i.e. ASV sequences longer than 260 bp or shorter than 250 bp). Taxonomy was assigned with the ‘assignTaxonomy’ function from the ‘dada2’ R package (v 1.12.1) and the SILVA (v. 132) reference database. ASVs sequences were then aligned using MAFFT (v.7.427) using the auto settings. A phylogentic tree was constructed from the resulting multiple sequence alignment with FastTree (v.2.1.11 Double Precision) using a generalized time-reversible model (‘-gtr’). The AVS table, taxonomy and tree were integrated using the ‘phyloseq’ R package (v.1.28.0). The ASV table was rarefied to 14932 counts per sample with vegan v2.5-6. Of 6056 sequenced samples, 24 had insufficient counts (<5000 counts per sample) and were excluded at the rarefaction stage. The final dataset thus contained 6032 samples and 22532 ASVs. Functional composition was inferred using PICRUSt2 (2.2.0b).

### Cell viability

Cell viability of was measured with CellTiter-Glo® Luminescent Cell Viability Assay (Promega) according to manufactures instructions. 10 size matched islets were used per replicate with 5 replicates per experiment. Luminescence was read with Promega GLOMAX^TM^ multi detection system.

### Statistical analysis

Data were checked for normality with the Shapiro–Wilk test. Paired or unpaired t-test was performed for normal continuous variables and the Wilcoxon signed rank test or Mann-Whitney for other variables. Spearman correlation was used for all correlation analysis. 2-way ANOVA with Šidák multiple comparison was used for glucose and insulin tolerance tests. Statistical analyses were performed using Prism, version 8.3.0 (GraphPad Software, US). Data are provided as mean with SEM . P-values < 0.05 were considered statistically significant. All authors had access to the study data and reviewed and approved the final manuscript.

## Supplementary information

**Table S1:** Patient characteristics of the Dutch participants of the HELIUS cohort (complementary to Figure 1A). Dutch origin people of the HELIUS cohort are shown. Data shown are mean ± SD for patient characteristics. Mann-Whitney test was used for statistical significance. Abbreviations: ND, no diabetes; T2D, Type 2 diabetes; HbA1c, glycated hemoglobin.

|  | ND | T2D | p value |
| --- | --- | --- | --- |
| n | 712 | 91 | n.d. |
| Sex (% female) | 57 | 34 | <0.0001 |
| Age (years) | 45.8 ± 13.0 | 61.7 ± 5.9 | <0.0001 |
| Body mass index (kg/m<sup>2</sup>) | 23.8 ± 3.5 | 29.2 ± 4.7 | <0.0001 |
| HbA1c (mmol/mol) | 34.3 ± 2.7 | 46.9 ± 8.2 | <0.0001 |

**Table S2:** Gut microbiota composition of HELIUS cohort (complementary to Figure 1A). Only a subset of Dutch origin people of the HELIUS cohort are shown. The 16S rRNA of the fecal microbiota was sequenced via miSeq (family level). Data shown are median. Abbreviations: ND, no diabetes; T2D, Type 2 diabetes

| Bacterial family | ND (n = 712) | T2D (n = 91) |
| --- | --- | --- |
| Bacteroidales_Rikenellaceae | 1.62 | 1.22 |
| Bacteroidales_Barnesiellaceae | 0.44 | 0.31 |
| Bacteroidales_Muribaculaceae | 0.54 | 0.67 |
| Bacteroidales_Prevotellaceae | 9.25 | 10.30 |
| Bacteroidales_Bacteroidaceae | 7.90 | 8.20 |
| Bacteroidales_Tannerellaceae | 0.71 | 0.78 |
| Clostridiales_Lachnospiraceae | 30.10 | 31.69 |
| Clostridiales_Peptostreptococcaceae | 1.08 | 0.71 |
| Verrucomicrobiales_Akkermansiaceae | 0.83 | 0.55 |
| Desulfovibrionales_Desulfovibrionaceae | 0.39 | 0.60 |
| Betaproteobacteriales_Burkholderiaceae | 0.68 | 0.74 |
| Enterobacteriales_Enterobacteriaceae | 0.35 | 1.08 |
| Bifidobacteriales_Bifidobacteriaceae | 2.34 | 1.57 |
| Coriobacteriales_Eggerthellaceae | 0.62 | 0.62 |
| Coriobacteriales_Coriobacteriaceae | 1.24 | 1.55 |
| Selenomonadales_Veillonellaceae | 1.74 | 1.92 |
| Selenomonadales_Acidaminococcaceae | 0.88 | 1.16 |
| Erysipelotrichales_Erysipelotrichaceae | 2.06 | 2.66 |
| Lactobacillales_Streptococcaceae | 0.52 | 0.96 |
| Clostridiales_Christensenellaceae | 1.95 | 1.68 |
| Clostridiales_Clostridiaceae_1 | 0.68 | 0.44 |
| Clostridiales_Ruminococcaceae | 30.35 | 27.28 |
| Other | 1.97 | 2.11 |

**Table S3.** Characteristics of selected participants from HELIUS cohort. Participants were randomly selected from the HELIUS cohort. People with Type 2 diabetes (T2D) were matched to controls without T2D according to age, sex and body mass index (BMI). Data shown are mean ± SD. Unpaired t-test was used for age and BMI. Mann Whitney test for fasting glucose and HbA1c. Abbreviations: n.d., not determined; BMI, body mass index; HbA1c, glycated hemoglobin.

|  | ND (n = 50) | T2D (n = 100) | p-value |
| --- | --- | --- | --- |
| Age (years) | 57.2 ± 6.8 | 56.8 ± 6.7 | 0.7440 |
| Female (%) | 62 | 62 | 1.0000 |
| BMI (kg/m<sup>2</sup>) | 29.7 ± 4.4 | 29.8 ± 5.0 | 0.9673 |
| Fasting glucose (mmol/L) | 5.35 ± 0.56 | 7.13 ± 1.85 | <0.0001 |
| HbA1c (mmols/mol) | 40.0 ± 4.2 | 54.5 ± 15.2 | <0.0001 |

**Table S4.**
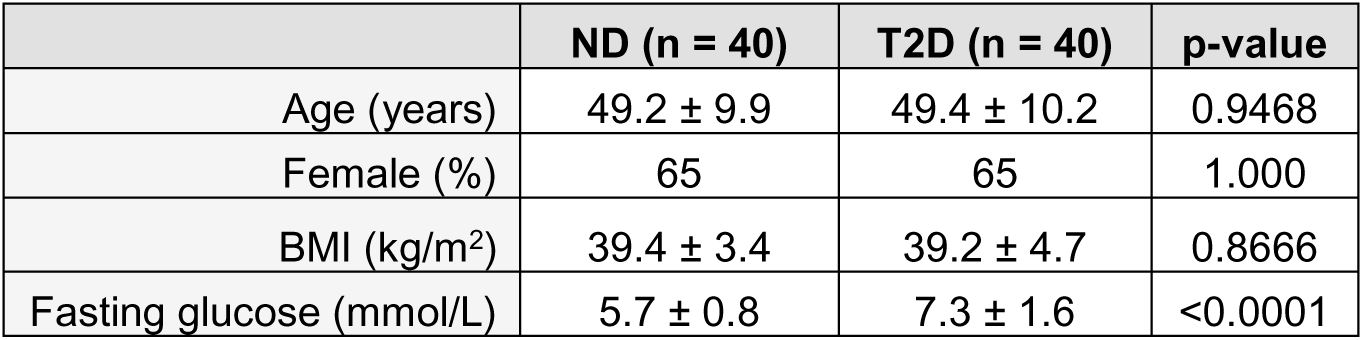
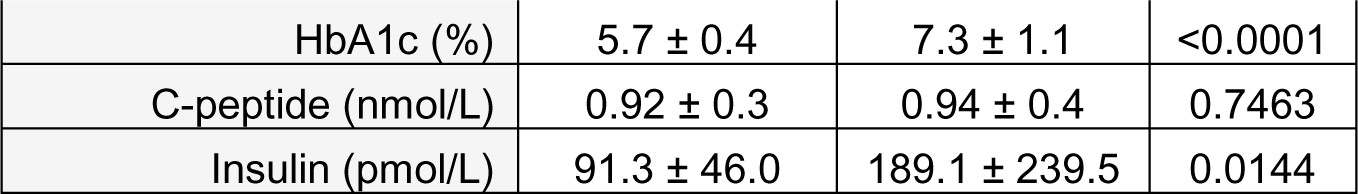
Characteristics of selected participants from BARIA cohort. Participants were randomly selected from BARIA cohort. People with Type 2 diabetes (T2D) were matched to controls without T2D according to age, sex and body mass index (BMI). Baseline samples were used before bariatric surgery. Data shown are mean ± SD. Unpaired t-test was used. Abbreviations: n.d., not determined; BMI, body mass index; HbA1c, Glycated hemoglobin; OD, optical density.

**Table S5.** Patient characteristics of diabetic individuals for human pancreatic biopsy analysis.

|  | **T2D (n = 5)** |
| --- | --- |
| Age (years) | 49.4 ± 10.2 |
| Sex (M/F) | 3/2 |
| BMI (kg/m<sup>2</sup>) | 27.1 ± 3.0 |
| HbA1c (%) | 7.8 ± 1.4 |
| Diabetes duration (years) | 9 ± 5 |

**Table S6.**
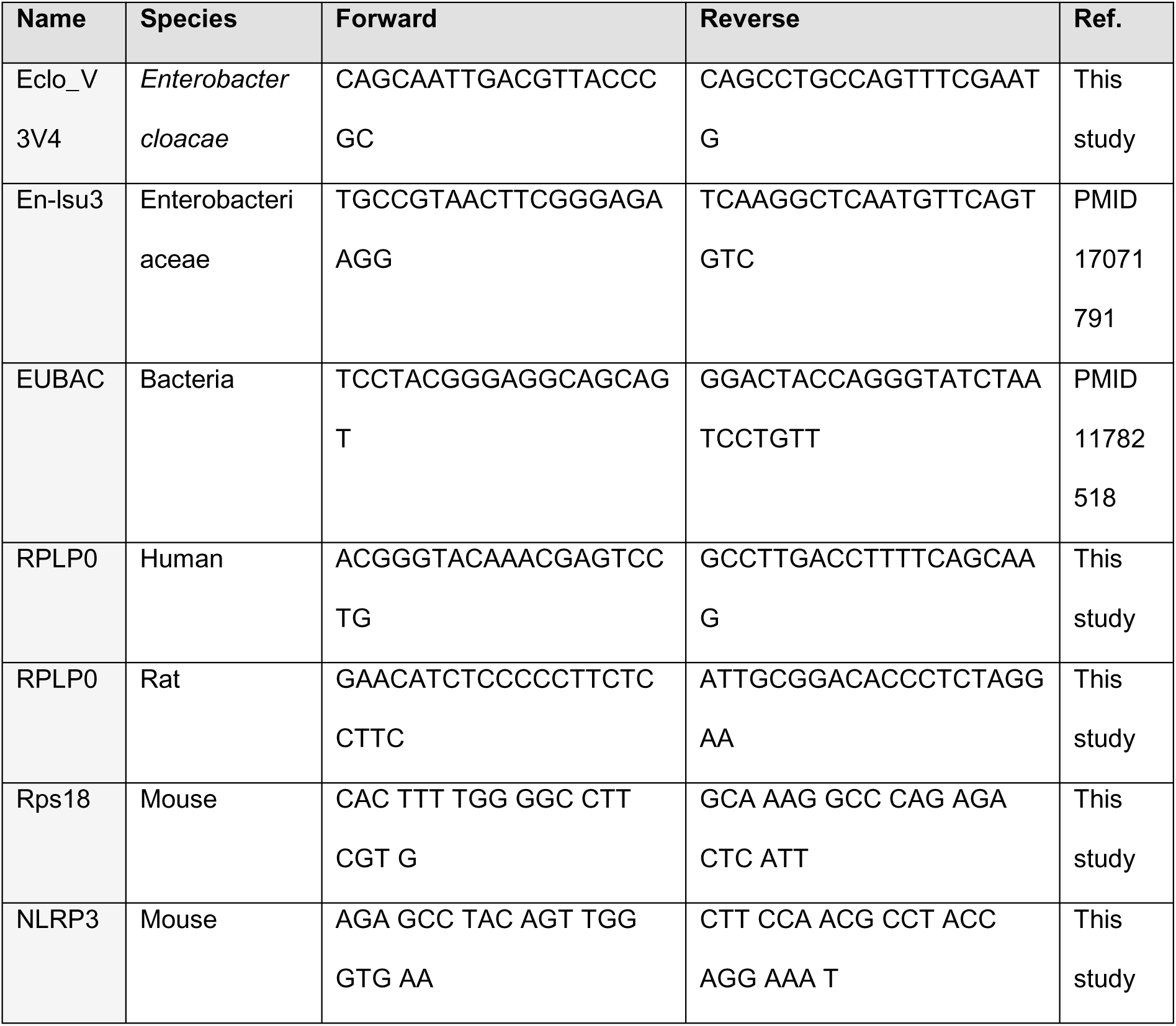
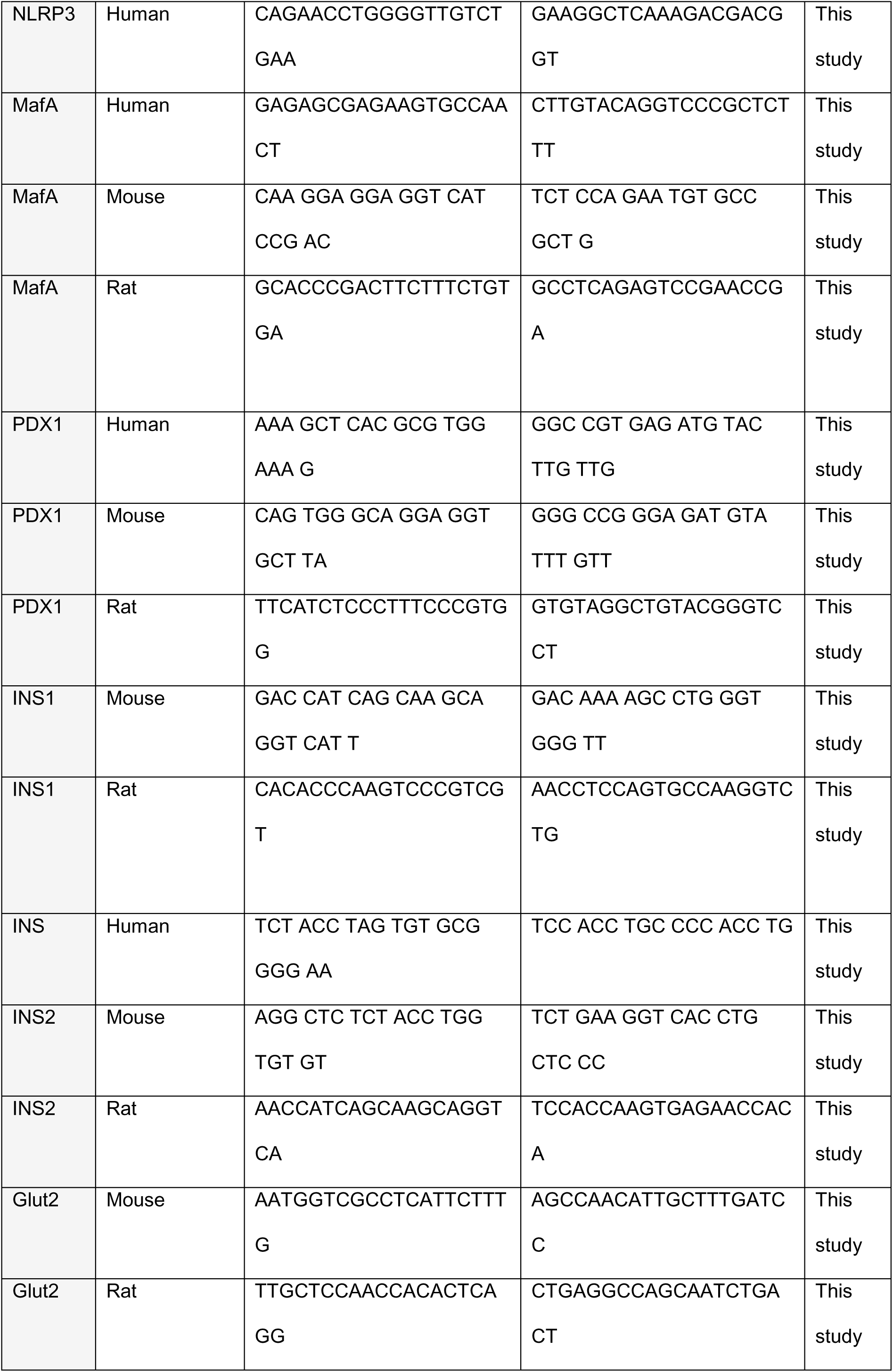
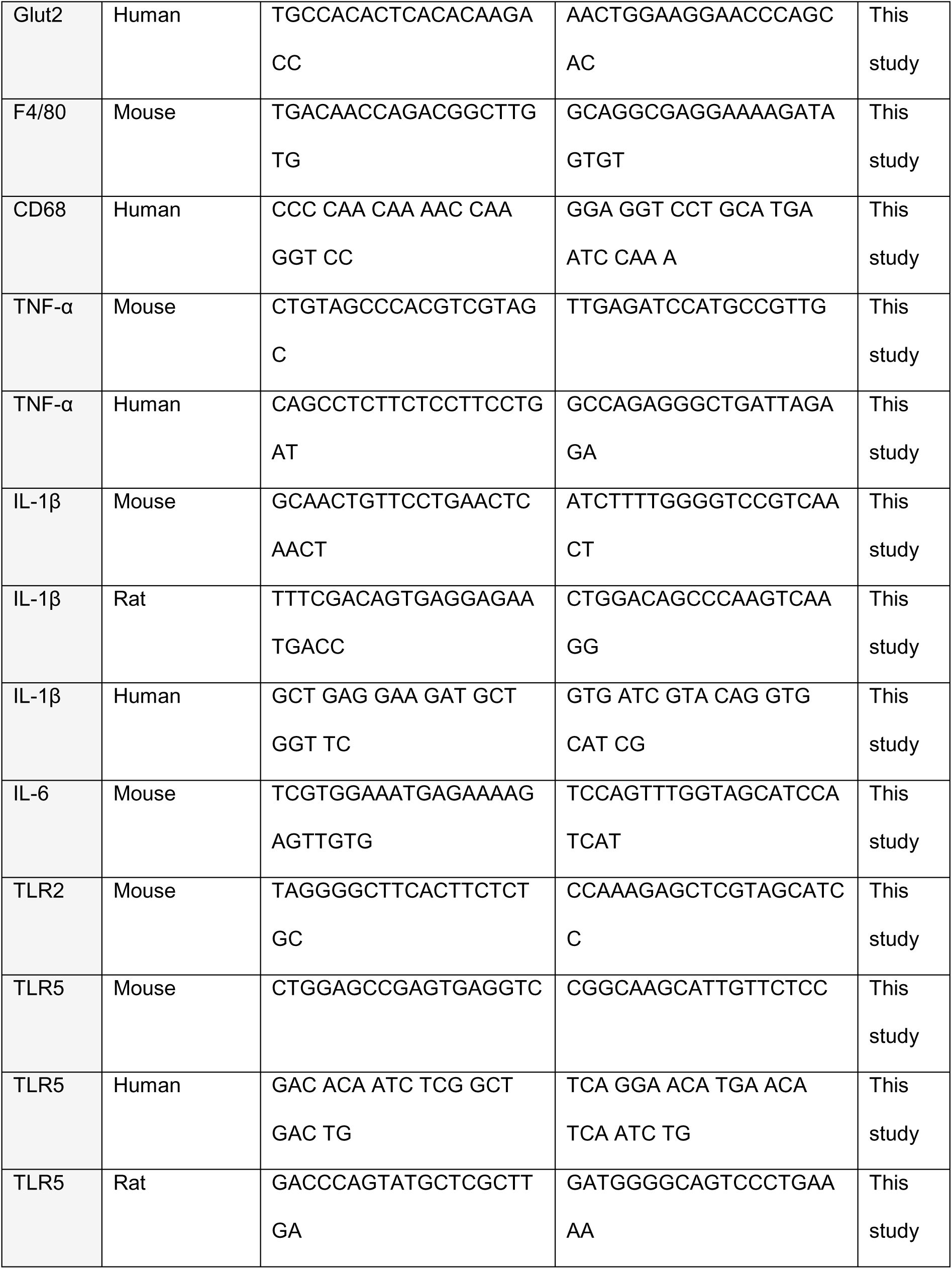
Primer sequences used in this manuscript (both in 5°3°direction).

---

**Figure S1.**
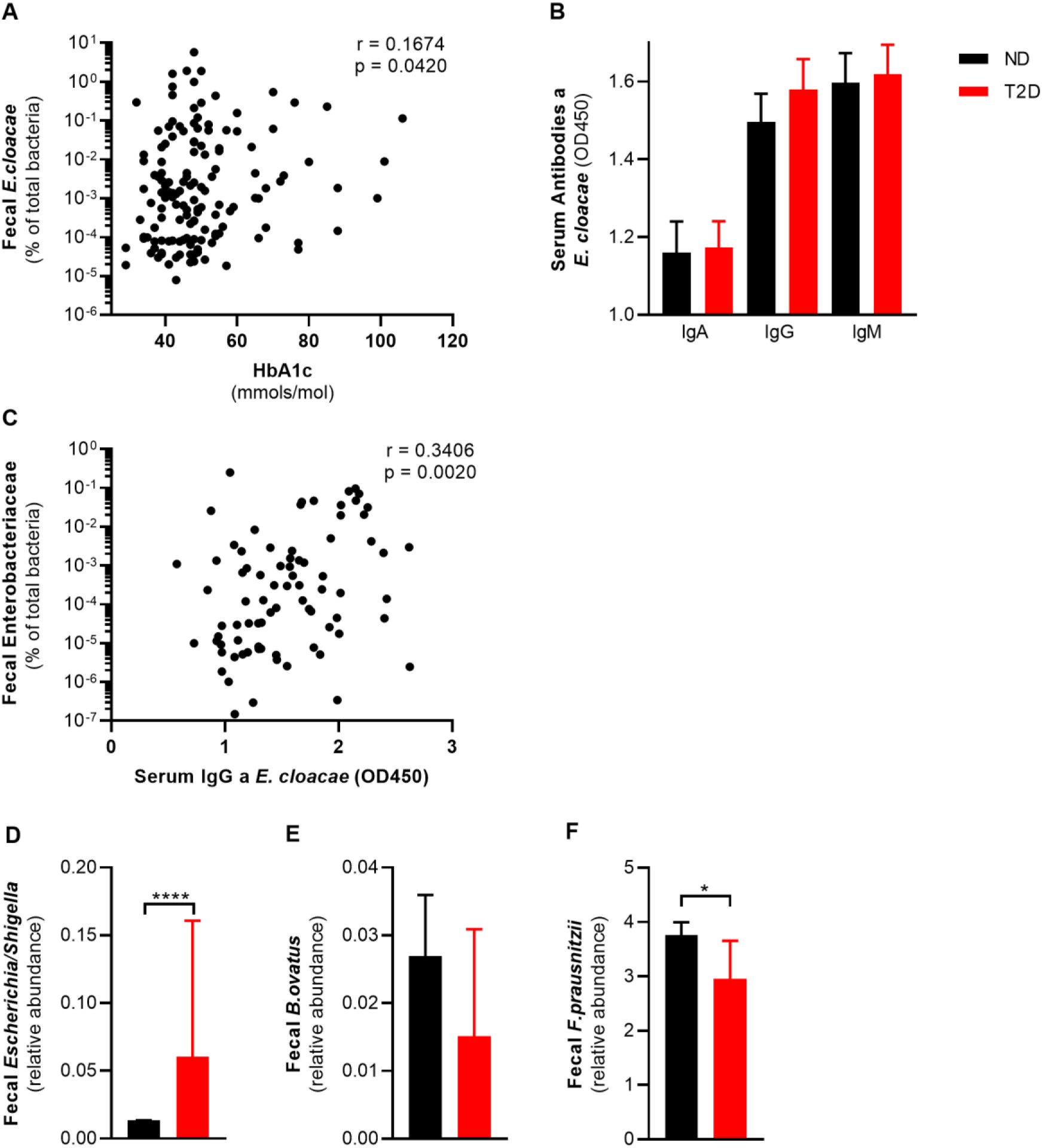
Fecal pathogens are associated with glucose intolerance. (A) Fecal *Enterobacter cloacae* positively correlates with glucose marker HbA1c in a subset of the HELIUS cohort (N = 150). (B) Serum antibodies against *E. cloacae* are not different between ND and T2D in subset of the HELIUS cohort (N = 80). (C) Serum IgG anti *E.cloacae* positively correlates with fecal Enterobacteriaceae (N = 80, Spearman correlation). (D) Fecal *Escherichia* is increased in T2D (HELIUS cohort, N = 803) (E) Fecal *Bacteroides ovatus* is non-significantly decreased in T2D (HELIUS cohort, N = 803) (F) Fecal *Fecalibacterium prausnitzii* is decreased in T2D (HELIUS cohort, N = 803) Data shown are mean ± SEM, except for gut microbiota (median with 95% confidence interval). Spearman correlation (A, C) and Mann Whitney test (D, F) was used: *p<0.05, ****p<0.0001. Abbreviations: ND, no diabetes; T2D, Type 2 diabetes; IgG, Immunoglobulin G; HbA1c, glycated hemoglobin.

**Figure S2.**
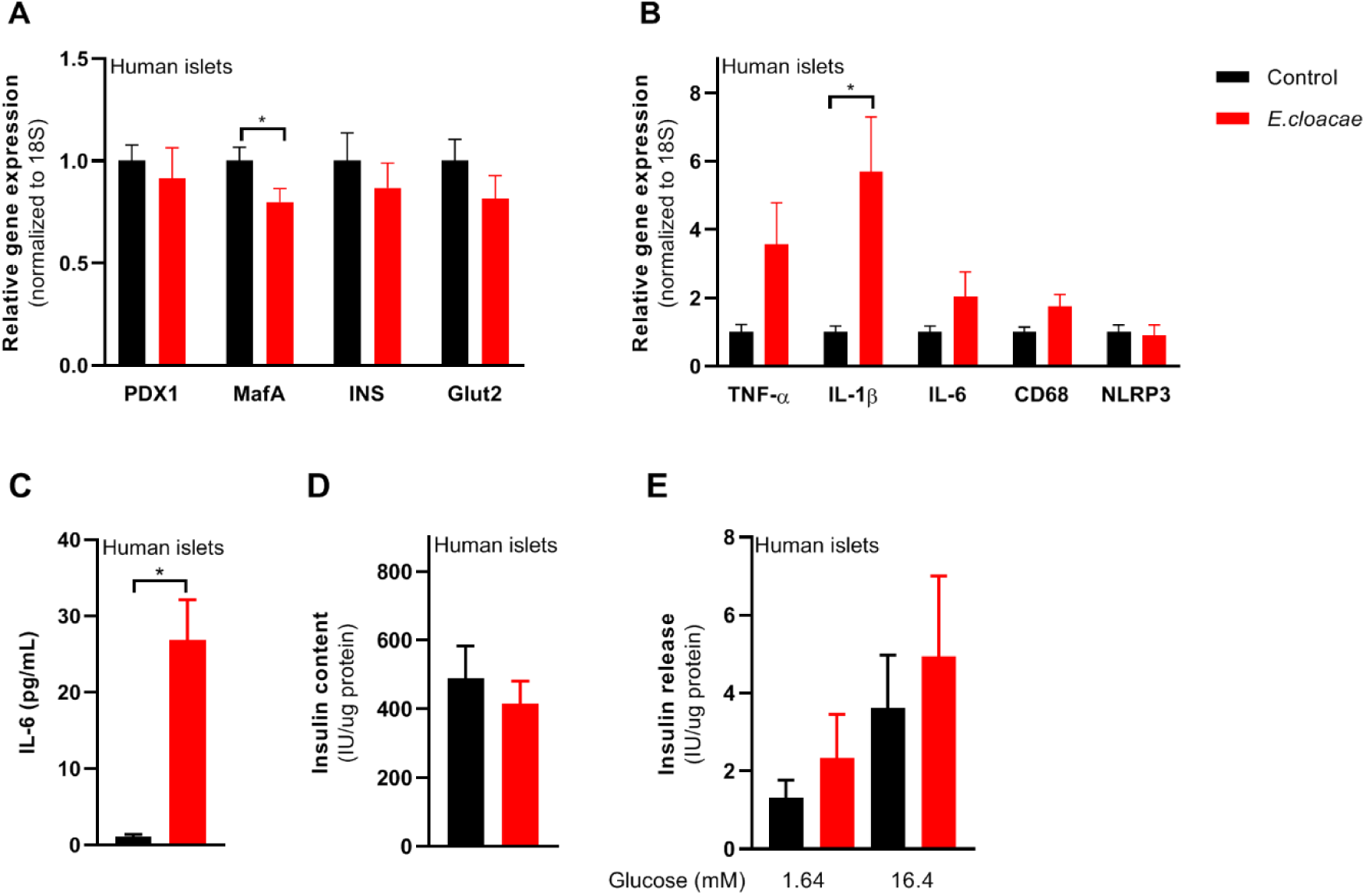
E.cloacae induces beta cell inflammation and partially dysfunction in human islets. Human islets were ordered from ProdoLabs and treated with *E.cloacae* (1E6 CFUs/mL) for 72h. (A) *E.cloacae* reduces beta cell marker expression in human islets. (B) *E.cloacae* induces beta cell inflammation in human islets. (C) *E.cloacae* induces IL-6 release from human islets. (D) *E.cloacae* slightly reduces insulin content in human islets. (E) *E.cloacae* slightly induces insulin hypersecretion in human islets. Data shown are mean ± SEM. Unpaired t-test was used for statistical analysis: *p<0.05, **p<0.01, ***p<0.001, ****p<0.0001. Abbreviations: INS1 and INS2, insulin 1 and 2; NLRP3, NACHT, LRR and PYD domains-containing protein 3; IL-1β, Interleukin 1 beta; IL-6, Interleukin 6; TLR, toll like receptor.

**Figure S3.**
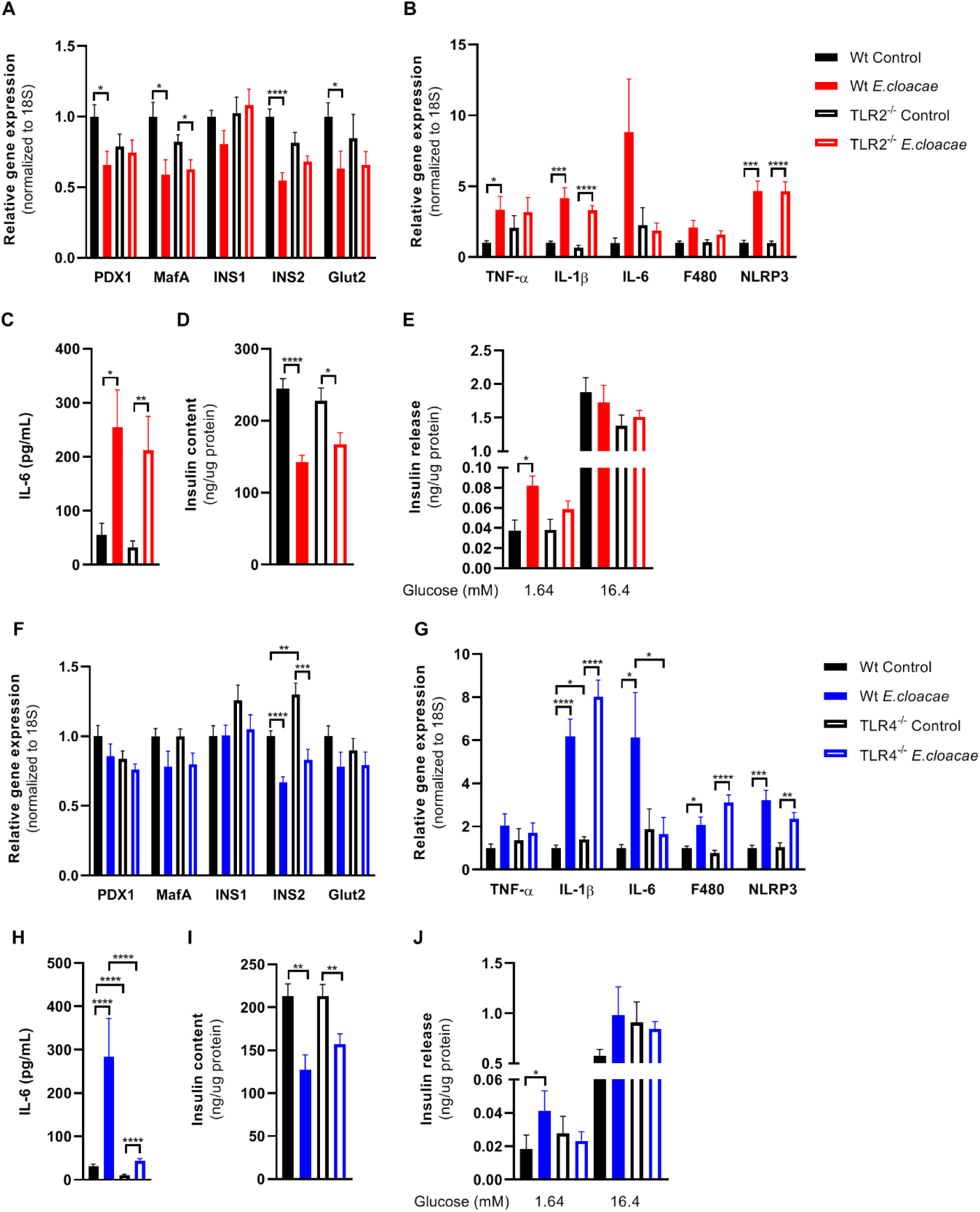
TLR2 and TLR4 knock out does not protect from beta cell inflammation and dysfunction. Freshly isolated pancreatic islets from C57BL6J TLR2^-/-^ (A-E) and TLR4^-/-^ (FJ) mice were treated with heat-inactivated *Enterobacter cloacae* (1E6 CFUs/mL) for 72h. (A) *E. cloacae* reduces expression of beta-cell genes both in wild-type and TLR2 knock out pancreatic islets of C57BL6J mice. (B) *E. cloacae* increases expression of inflammatory genes in wild-type and TLR2 knock out pancreatic islets of C57BL6J mice. (C) *E. cloacae* increases secreted IL-6 from by wild-type and TLR2 knock out pancreatic islets of C57BL6J mice. (D) *E. cloacae* reduces insulin content in wild-type and TLR2 knock out pancreatic islets of C57BL6J mice. (E) *E. cloacae* induces insulin hypersecretion in wild-type and TLR2 knock out pancreatic islets during low-glucose concentrations of C57BL6J mice. (F) *E. cloacae* reduces expression of beta-cell genes both in wild-type and TLR4 knock out pancreatic islets of C57BL6J mice. (G) *E. cloacae* increase expression of inflammatory genes in islets in wild-type and TLR4 knock out pancreatic islets of C57BL6J mice. (H) *E. cloacae* increases secreted IL-6 from by wild-type and TLR4 knock out pancreatic islets of C57BL6J mice. (I) *E. cloacae* reduces insulin content in wild-type and TLR4 knock out pancreatic islets of C57BL6J mice. (J) *E. cloacae* induces insulin hypersecretion at low glucose concentrations in wild-type and TLR4 knock out pancreatic islets of C57BL6J mice. Data shown are mean ± SEM. Unpaired t-test (A-C, F-H) or Mann-Whitney test (D, E, I, J) was used for statistical analysis (mean ± SEM, 3 representative experiments per panel). Abbreviations: PDX1, pancreatic and duodenal homeobox 1; INS1 and INS2, insulin 1 and 2; NLRP3, NACHT, LRR and PYD domains-containing protein 3; TNF-α, tumor necrosis factor-alpha; IL-1β, Interleukin 1 beta; IL-6, Interleukin 6; TLR2, Toll-like receptor 2.

**Figure S4.**
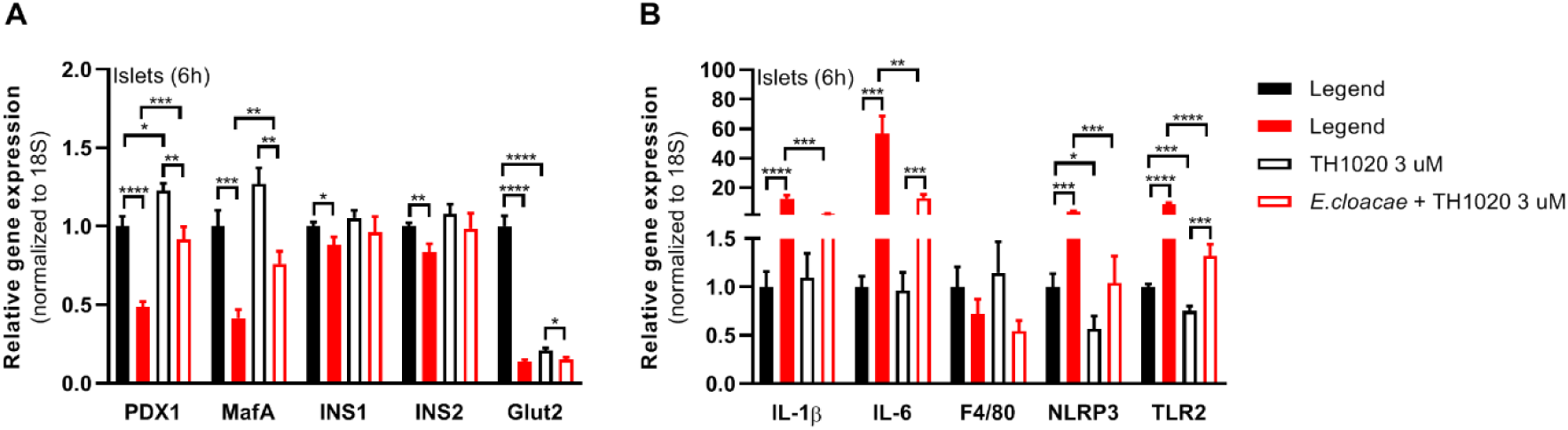
TLR5 inhibitor reduces bacteria induced beta cell dysfunction. Pancreatic islets were isolated from C57BL6J mice and incubated with TLR5 inhibitor (3 uM) and *E.cloacae* (1E6 CFUs/mL) for 6h. (A) TLR5 inhibitor TH1020 reverses bacteria induced pancreatic islet dysfunction. (B) TLR5 inhibitor TH1020 reduces bacteria induced pancreatic islet inflammation. Data shown are mean ± SEM. Unpaired t-test was used for statistical analysis: *p<0.05, **p<0.01, ***p<0.001, ****p<0.0001. Abbreviations: INS1 and INS2, insulin 1 and 2; NLRP3, NACHT, LRR and PYD domains-containing protein 3; IL-1β, Interleukin 1 beta; IL-6, Interleukin 6; TLR, toll like receptor.

**Figure S5.**
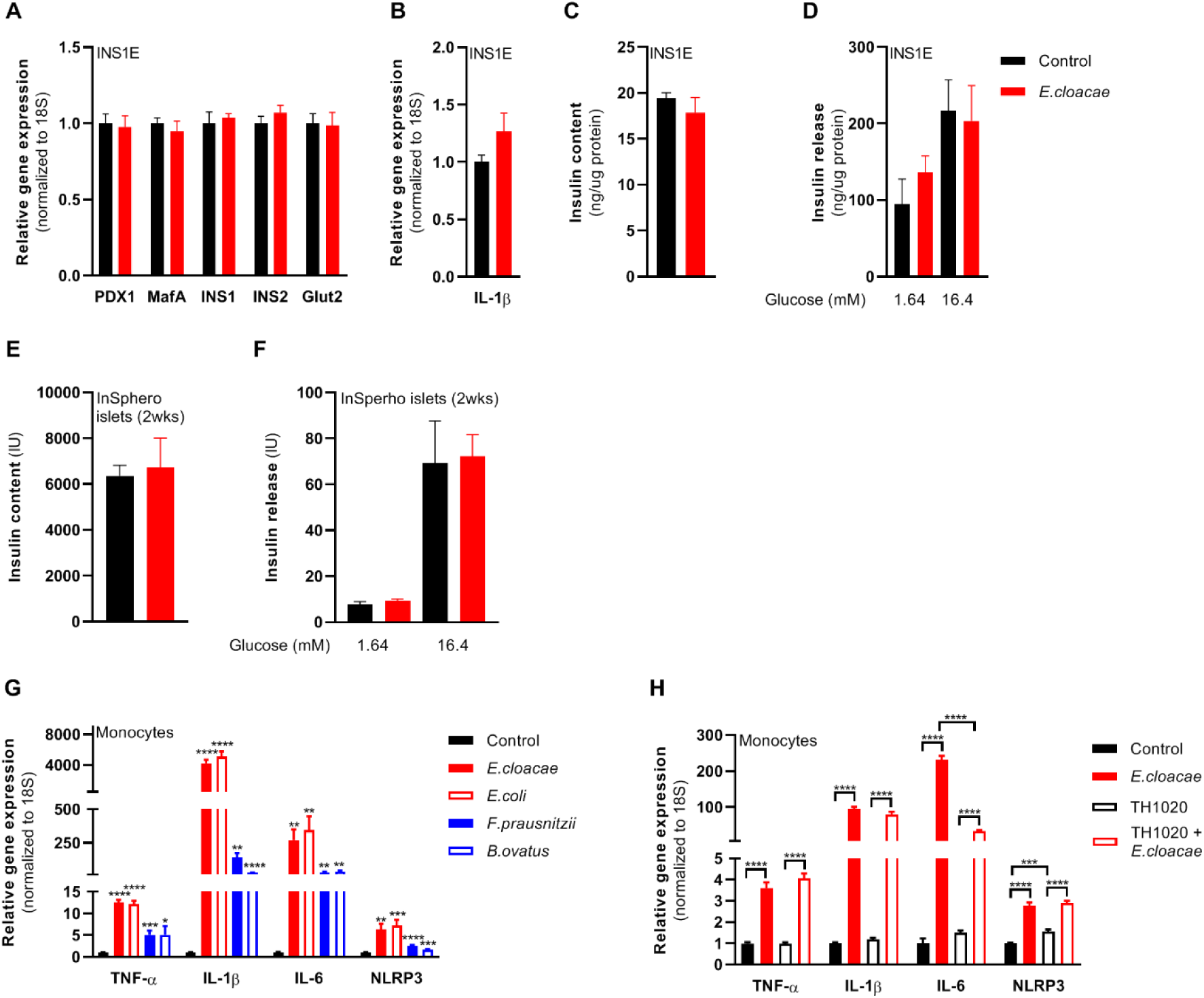
Pure beta cells do no respond to bacteria, but monocytes. Pure beta cells (INS1E, A-D) and modified human islets without immune cells (InSphero islets, E-F) were treated with *E.cloacae* for 72h. Human monocytes were treated with bacteria (1E6 CFUs/mL) or TLR5 inhibitor TH1020 (3 uM) for 24h (G-H). (A) *E.cloacae* does not reduce beta cell marker expression. (B) *E.cloacae* does not induce beta cell inflammation. (C) *E.cloacae* does not reduce insulin content in beta cells. (D) *E.cloacae* does not induce insulin hypersecretion in beta cells. (E) *E.cloacae* does not reduce insulin content in InSphero islets. (F) *E.cloacae* does not induce insulin hypersecretion in InSphero islets. (G) Opportunistic pathogens induce more inflammation than beneficial bacteria in human monocytes. (H) TLR5 inhibitor TH1020 reduces IL-6 expression in *E.cloacae* treated monocytes.. Data shown are mean ± SEM. Unpaired t-test was used for statistical analysis: *p<0.05, **p<0.01, ***p<0.001, ****p<0.0001. Abbreviations: INS1 and INS2, insulin 1 and 2; NLRP3, NACHT, LRR and PYD domains-containing protein 3; IL-1β, Interleukin 1 beta; IL-6, Interleukin 6; TLR, toll like receptor.

**Figure S6.**
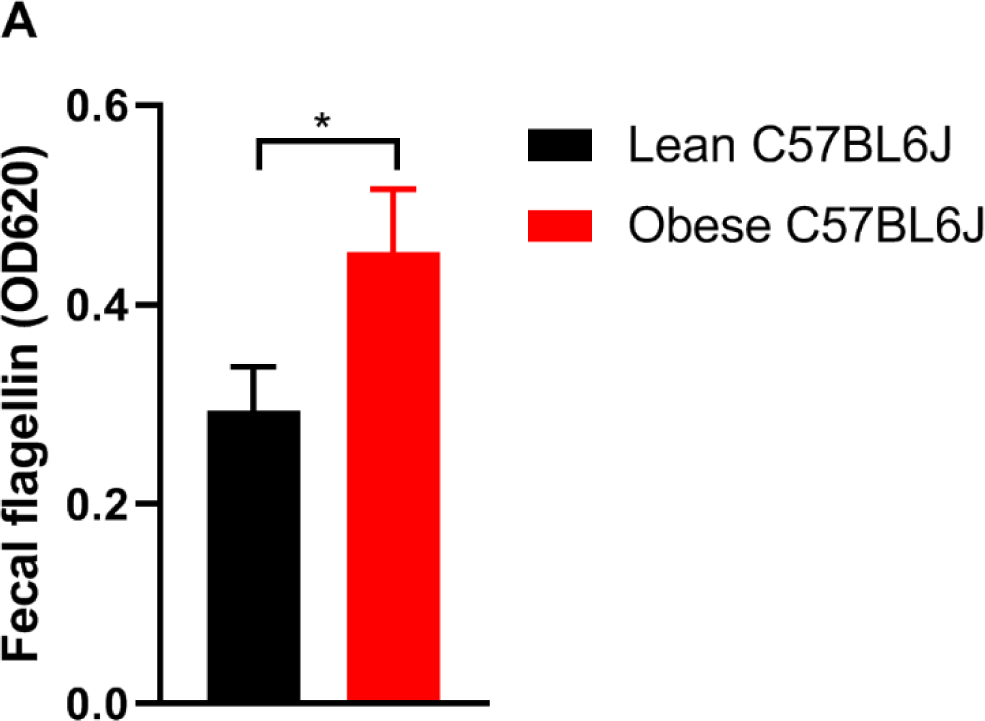
High fat diet feeding increases fecal flagellin content in mice. C57BL6J mice were on a high fat diet (60% kcal fat) for 12 weeks. Fecal flagellin was measured in homogenized fecal samples with HEK TLR5 reporter cells.

**Figure S7.**
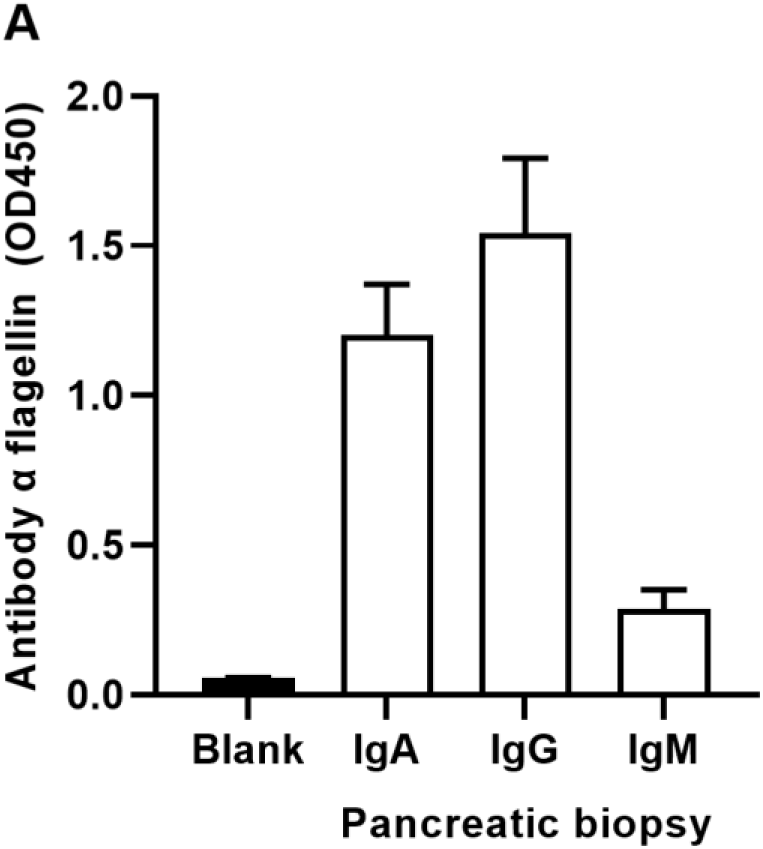
Human pancreatic biopsies harbor antibodies against bacterial flagellin. Antibodies against flagellin were measured in homogenates of pancreatic biopsies from people with T2D undergoing pancreatic surgery.

**Figure S8.**
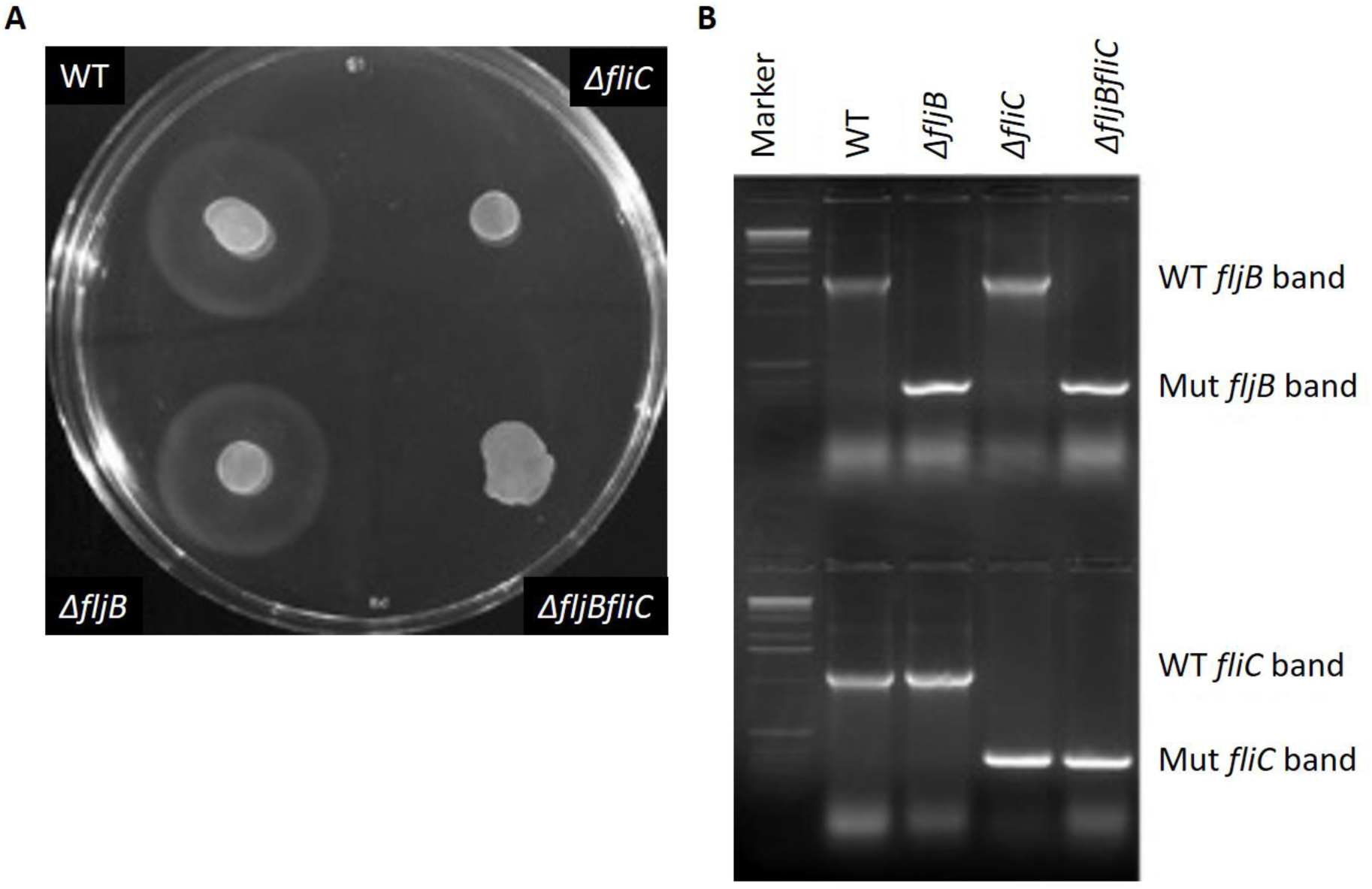
The flagellin gene was deleted in Enterobacter cloacae. (A) Motility assay and (B) PCR control showing that *E.cloacae* does not express flagella.

